# Brain signatures of semantic activation for words that do not exist

**DOI:** 10.64898/2026.04.29.721646

**Authors:** Rolando Bonandrini, Simona Amenta, Simone Sulpizio, Gianpaolo Basso, Marco Marelli, Marco Tettamanti

## Abstract

The experience of making sense of novel words is ubiquitous in human communication. Still, novel words have traditionally been considered as meaningless by cognitive research. Here we combined behavioral, univariate and multivariate fMRI techniques, and computational modelling to explore whether and how novel words activate the neurocognitive hallmarks of semantic processing triggered by existing words, and whether this process is influenced by the presence of familiar functional linguistic elements, i.e., morphemes. We observed that semantic activation for novel words is comparable to that of existing words, provided that novel words contain a concatenation of identifiable morphemes. In addition, representational similarity analysis highlighted that existing words and novel words containing morphemes (but not novel words not containing morphemes) can activate fine-grained semantic representations. These results suggest that the difference in the neurocognitive underpinnings of semantic processing for existing and novel words might be quantitative rather than qualitative and based on how reliably linguistic form points to meaning.

---

Words are the main currency in human information exchange. An average 20-year-old native speaker of American English knows around 42000 words^1^, which can be used every day in speech, reading and writing to convey and retrieve meaning. Still, we learn a new word every two days^1^ and there are always new ones to learn, as every language has virtually infinite potential for expansion^2, 3^. There is indeed a gap between the words we know and the words that may potentially exist which are still unknown to us. If so, then what is a word “in us” (i.e., which neurocognitive mechanisms does it activate) if it is not yet a word “to us”? Is there some kind of “brick wall” inside our head that—from the standpoint of a human receiver—divides existing words from novel conglomerates of letters?

Localizing and outlining the organization of linguistic processing and how it relates to what we know about the world (i.e., semantics) are critical challenges in cognitive science and neuroscience. Since as early as in the late 19^th^ century, an array of cerebral areas mapping the so-called “word treasury” have been described. These areas mainly involve left inferotemporal, inferior frontal and parietal cortices for written language processing, superior temporal and inferior frontal cortices for spoken language, superior prefrontal and posterior parietal cortices for meaning extraction, and anterior temporal areas acting as a semantic hub (i.e., as a mediator of interactions between modality-specific sources of information)^4–18^.From a cognitive standpoint, the architecture of word knowledge has been traditionally conceptualized as a “lexicon”, a “dictionary” of the mind containing units that, once activated, trigger the retrieval of the corresponding meaning in the so-called “semantic system”^19–24^. However, the borders between the lexicon and the semantic system are actually rather nuanced. Meaning can indeed be activated even in absence of word recognition (e.g., through activation of orthographically similar “neighbor” words)^25, 26^. In addition, there seems to be no one-to-one mapping between the lexicon and the brain, in that brain areas related to lexical processing are also involved in other functions^27^, and—conversely—multiple areas may contribute to what is commonly labelled “lexical access”^28, 29^. Finally, there seem to be no well-defined anatomical boundaries between the neural correlates of the lexicon and those of semantics^30–32^.

Such blurred lines between the lexicon and the semantic system suggest that word knowledge is neurocognitively fleeting and elusive in nature^33, 34^, despite the researchers’ best efforts to encage it in a “dictionary”.

One may even wonder to what extent words represent a psychological reality at all, as different standpoints exist in the literature in this regard. Earlier formulations proposed that words are a fundamental representational unit: each known word would be stored as a distinct and independent representation in the human mind advocating a localist organization of lexical and semantic knowledge proposed a distinct and independent representation in the human mind for each individual known word^19–23^. Psycholinguistic research on morphemes added nuance to this picture: statistically relevant units of form contained in words (e.g., *over*-, *-simply-*, and –*ify* in *oversimplify,* representing a prefix, a stem and an affix, respectively) can facilitate or hamper lexical processing/word recognition, thus suggesting that lexical-semantic mechanisms do not necessarily require representation of full words in a mental dictionary^35–43^. An even more radical approach to lexical-semantic processing is offered by distributed frameworks, according to which each given word is encoded via multiple units and the same units are associated to multiple words (as anticipated by the connectionist tradition^44–46^). These latter accounts suggest that assuming the existence of a “mental dictionary” might be unnecessary: computational data revealed that the degree of activation of semantic representations in a distributed system that statistically connects meaning to form is indeed able to account for word recognition effects even without assuming the existence of a lexicon altogether^34, 47^.

It appears therefore evident that qualifying the relation between word form (i.e., orthography/phonology) and meaning is critical for understanding how lexical-semantic knowledge is represented in the human mind and brain. Indeed, meta-analytical work on patients with chronic stroke aphasia highlighted that the two factors that most prominently explain variance in aphasic deficits are those related to linguistic form and meaning^48, 49^. Recent innovations in computational modelling have made it possible to gauge form-meaning mapping, by highlighting the role of morphemes and sublexical units of variable size (i.e., *ngrams*: bigrams, trigrams, etc.) in acting as interfaces towards semantics^50–56^. Such techniques have offered a computational operationalization of how form-meaning associations at the sub-word level may be leveraged to extract meaning not only from words we know, but also from words we do not know as well^57–60^. For known words, this process entails merging semantic information coming from the word itself with that coming from the sublexical units that compose it^54, 55, 61, 62^. For novel words, *ngram*-based form-meaning mapping represents the foundational mechanism to extract meaning on top of which morphological information is leveraged, whenever available^63^.

The unprecedented opportunity to gauge meaning extraction from novel words via computational techniques brings critical theoretical implications related to the study of the neurocognitive underpinnings of language processing. Indeed, for a relevant portion of the literature that adopted the “mental dictionary” metaphor to conceptualize lexical-semantic processing, if a word is not in the dictionary (as in the case—by definition—of novel words), it cannot trigger the cascade of events that culminates with semantic access^23, 64^. Accordingly, a novel word could—at most—be read by converting its orthographic form into a phonological one by adopting language-specific rules. The possibility to extract meaning from novel words suggests instead that reading words that are not part of the “dictionary” is not entirely resolved by sublexical grapheme-phoneme conversion mechanisms. Moreover, it indicates that words are *per se* not necessary for semantic activation during language processing^58, 59^: semantic activation can follow domain-general processes that allow meaning extraction from the form of a linguistic stimulus, regardless of whether such stimulus is an existing word or not.

In this context, the emerging scenario is one whereby words are the tip of an iceberg of form-meaning associations: they just happen to represent the most reliable way to access semantics; yet meaning can also be retrieved (with progressively decreasing efficiency) from morphemes^65, 66^ and *ngrams*^54, 61^. Depending on which gateway can be relied upon to access meaning (i.e., the word itself, morphemes, *ngrams*), letter strings may theoretically be positioned along a continuum encompassing complex existing words (that contain both the word itself and constituent morphemes as cues to meaning; e.g., “*pianist*”), simple existing words (for which a semantic representation is available, but no morphemes are embedded in the string; e.g., “*table*”), complex novel words (novel words for which morphemes can help access meaning, e.g., “*windowist*”), and simple novel words (e.g., “*mimpas*”; for which access to meaning only depends on *ngrams*). This framework presumes the existence of quantitative differences between words and novel words, with potential overlapping substrates underlying their semantic processing.

Still, whether novel words are able to activate semantic representations (and their underlying neural substrates) to a similar extent and in a similar way as existing words remains unclear. Here we combined behavioral, univariate and multivariate fMRI techniques, and computational modelling to probe the existence of a gradient in semantic activation between existing and novel words, while modulating Lexicality (i.e., whether a word is existing or novel) and Morphological complexity (i.e., “simple”: absence of morphemes vs. “complex”: presence of morphemes). More specifically, in behavioral and univariate fMRI data we explored similarities and differences between existing and novel words in how they engage the neurocognitive underpinnings of semantic processing, as well as the possible intervening role of morphemes in the activation of the semantic system. Furthermore, we adopted a computational model derived from distributional semantics to extract high-dimensional semantic representations induced by existing and novel words. We performed Representational Similarity Analysis (RSA) on multivariate fMRI data to explore the extent to which the brain responds to existing and novel words in a way that is aligned with such fine-grained representations corresponding to points in a high-dimensional semantic space.

Results revealed that novel words can activate the same neurocognitive hallmarks of semantic processing as those triggered by existing words—as long as they contain morphemes. In addition, representational similarity analyses highlight that existing words and novel words containing morphemes can activate fine-grained semantic representations which can be gauged by a distributional computational model.

## Results

### Behavioral results

During 3T fMRI scanning (**Fig.1c**), 32 healthy participants were presented with 192 written Italian verbal stimuli (**Supplementary Table S1**, **Fig.1a**). Stimuli were modulated according to the variables “Lexicality” (existing word vs. novel word) and “Morphological complexity” (simple: absence of morphemes vs. complex: presence of morphemes), with a total of 48 stimuli per condition. During the presentation of each stimulus, participants were instructed to reflect upon the meaning it evoked, so as to be able to provide (after a blank screen) semantic ratings jointly encompassing imageability and valence by positioning a cursor in a two-dimensional grid (**Fig.1b**).

We first made sure that behavioral responses (**Fig.2a**) were not converging towards the center of the axes (i.e., null hypothesis: participants’ ratings are not different from zero). This proved to be the case for all categories of stimuli (morphologically complex existing words *t*(51.61)= 25.09, *p*< 0.001; morphologically simple existing words *t*(60.66)= 30.90, *p*< 0.001; morphologically complex novel words *t*(35.95)= 20.98, *p*< 0.001; morphologically simple novel words *t*(36.20)= 34.52, *p*< 0.001).

We then explored the effects of Lexicality, Morphological complexity, and their interaction on imageability and valence ratings, as well as on a composite behavioral metric of semantic activation that we labelled as “Overall Semantic Loading” (OSL; see the Methods section for further details; see also **Fig. 1e**), in which zero represents the absence of semantic activation.

**Fig. 1.**
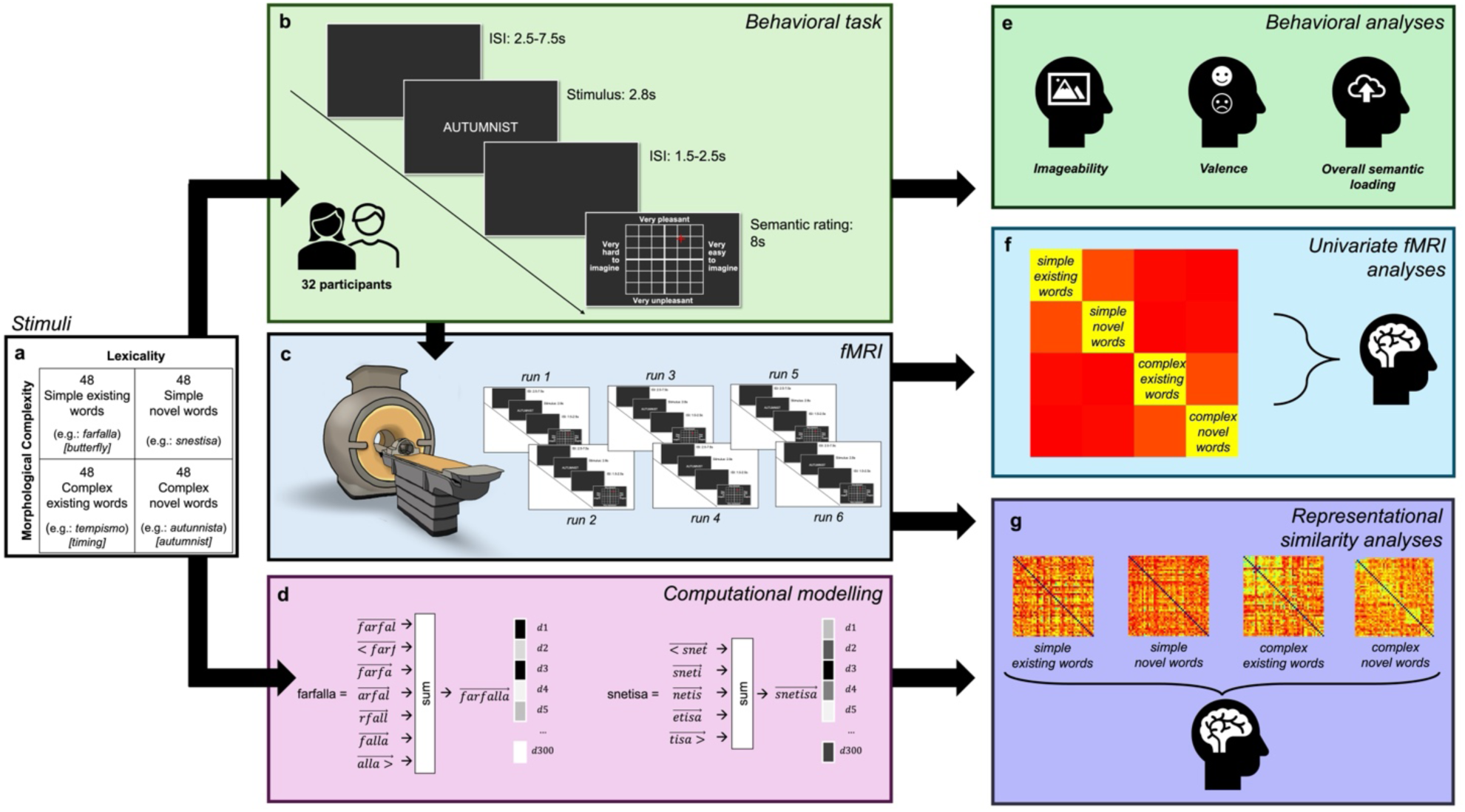
| Methodological outline of the study. (**a**) A total of 192 verbal stimuli were used, which resulted from the joint manipulation of Lexicality and Morphological Complexity. (**b**) Participants were asked to read the stimuli and provide, on a two-dimensional grid, ratings of imageability and valence. ISI: inter-stimulus interval. (**c**) Functional MRI scanning was carried out throughout the task. The 2s of stimulus presentation represented the event of interest for subsequent analyses. (**d**) We used FastText^54^ to induce vector representations for existing and novel words. (**e**) In behavioral analyses we explored the effects of Lexicality and Morphological complexity on Imageability and Valence ratings, as well as on a pooled index, i.e., “Overall Semantic Loading”. (**f**) In Univariate fMRI analyses we tested the effects of Lexicality and Morphological complexity on the neural response in the whole brain. (**g**) A multivariate Representational Similarity Analysis was carried out to explore (in the different categories of stimuli) alignment between brain data and the fine-grained vector representations of meaning extracted from FastText.

**Fig. 2.**
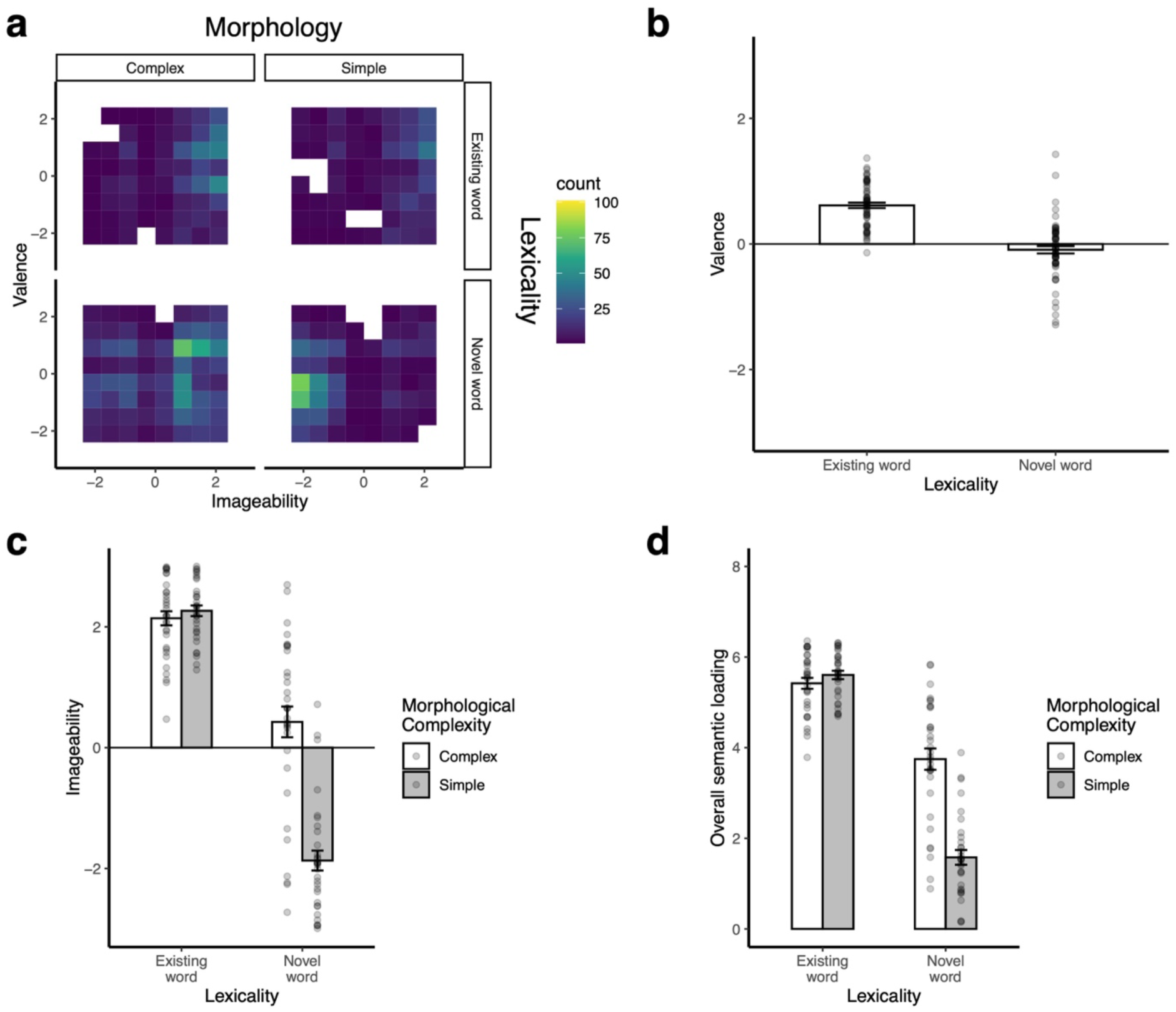
| Behavioral data. (**a**) Distribution of valence and imageability ratings from participants across the four categories of stimuli. (**b**) Lexicality effect on valence ratings. (**c**) Lexicality-by-Morphological complexity interaction in imageability ratings. (**d**) Lexicality-by-Morphological complexity interaction in OSL. OSL: Overall Semantic Loading. Error bars indicate standard error. Points depict scores from individual participants.

We observed a main effect of Lexicality in valence ratings (*F*(1,188)= 20.459, *p*< 0.001), in that existing words were rated more positively than novel words overall (**Fig.2b**). As for imageability, we detected a significant main effect of Morphological complexity (*F*(1, 187.97)= 181.72, *p*< 0.001), a significant main effect of Lexicality (*F*(1, 187.97)= 1318.64, *p*< 0.001), and a significant lexicality-by-Morphological complexity interaction (*F*(1, 187.97)= 225.64, *p*< 0.001), whereby all categories of stimuli were different from one another (min|*t*(188)|= 15.06, *p_Bonferroni-corrected_*<0.001), **Fig.2c**), with the only exception of morphologically simple existing words and morphologically complex existing words (t(188)= –1.090, *p_Bonferroni-corrected_* =1.000). Results on valence and imageability were recapitulated by OSL: analyses on OSL revealed a significant main effect of Morphological complexity (*F*(1,188)= 171.63, *p*< 0.001), a significant main effect of Lexicality (*F*(1,188)= 1415.61, *p*< 0.001), and a significant Lexicality-by-Morphological complexity interaction (*F*(1,188)= 241.55, *p*< 0.001). Post-hoc pairwise comparisons revealed that all categories of stimuli were different from one another (min|*t*(188)|= 15.61, *p_Bonferroni-corrected_*<0.001, **Fig.2d**), with the only exception of morphologically simple existing words and morphologically complex existing words (*t*(188)=-1.726, *p*=0.516). More specifically, morphologically simple novel words were associated with the lowest semantic activation, existing words (regardless of their morphological complexity) had maximal semantic activation, whereas morphologically complex novel words showed an intermediate degree of semantic activation. The OSL metric turned out to be significantly greater than zero in all conditions (morphologically complex existing words *t*(54.86)= 37.72, *p*< 0.001; morphologically simple existing words *t*(72.27)= 42.33, *p*< 0.001; morphologically complex novel words *t*(35.08)= 15.40, *p*< 0.001; morphologically simple novel words *t*(36.22)= 9.28, *p*< 0.001), which suggests that all types of stimuli yielded some degree of semantic activation.

### Univariate fMRI results

Univariate fMRI analyses (**Fig.1f**) were conducted to explore whether the brain’s semantic network^16^ is activated while processing novel words and whether this effect is modulated by the availability of morphemes.

We modelled the Blood-Oxygenation-Level-Dependent (BOLD) response recorded during stimulus presentation within each trial. In analyses on individual participant data we extracted simple-effects estimates for each of the four stimuli categories of interest (i.e., weight=1 for each condition of interest, 0 otherwise), which were then brought to a group level analysis where they were modelled according to a full-factorial design to instantiate a 2×2 (Lexicality by Morphological complexity) structure. We then ascertained that the effects of Lexicality, Morphological complexity and their interaction involved brain areas compatible with the semantic system (**Supplementary Table S2**). This was true (% overlapping voxels > 0) for all clusters of significant activation (peak-level threshold: p<0.05 Family-Wise-Error corrected, minimum cluster size= 30 voxels) apart from a cluster in the middle occipital gyri (Lexicality effect), and one in the cerebellum (Lexicality-by-Morphological complexity interaction). Still, it is worth noting that there exist recent evidence pointing to the involvement of cerebellar structures in semantics^67–70^. The results of our univariate analyses, as far as the Lexicality factor is concerned, replicate previous findings in the neuroimaging literature^11, 12, 71^ (**Supplementary Table S3**). In particular, the left angular gyrus, the left precuneus, the left superior frontal gyrus, the left middle temporal gyrus, and the right supramarginal gyrus turned out to be more strongly recruited by processing of existing words (as opposed to novel words; **Fig. 3a**, **Supplementary Table S4**). Conversely, the opercular portion of the left inferior frontal gyrus and both the left and right middle occipital gyri were more activated during processing of novel words than during the processing of existing words (**Fig. 3b**, **Supplementary Table S5**). Consistently with previous evidence^72, 73^, the main effect of Morphological complexity involved a similar set of fronto-parietal areas (**Supplementary Table S6**): the left supramarginal and middle occipital gyri were more activated for morphologically simple than for morphologically complex stimuli (**Fig. 3c**, **Supplementary Table S7**), whereas the opposite trend was detected in the left angular gyrus, left precuneus, and left superior frontal gyrus (**Fig. 3d**, **Supplementary Table S8**).

**Fig. 3.**
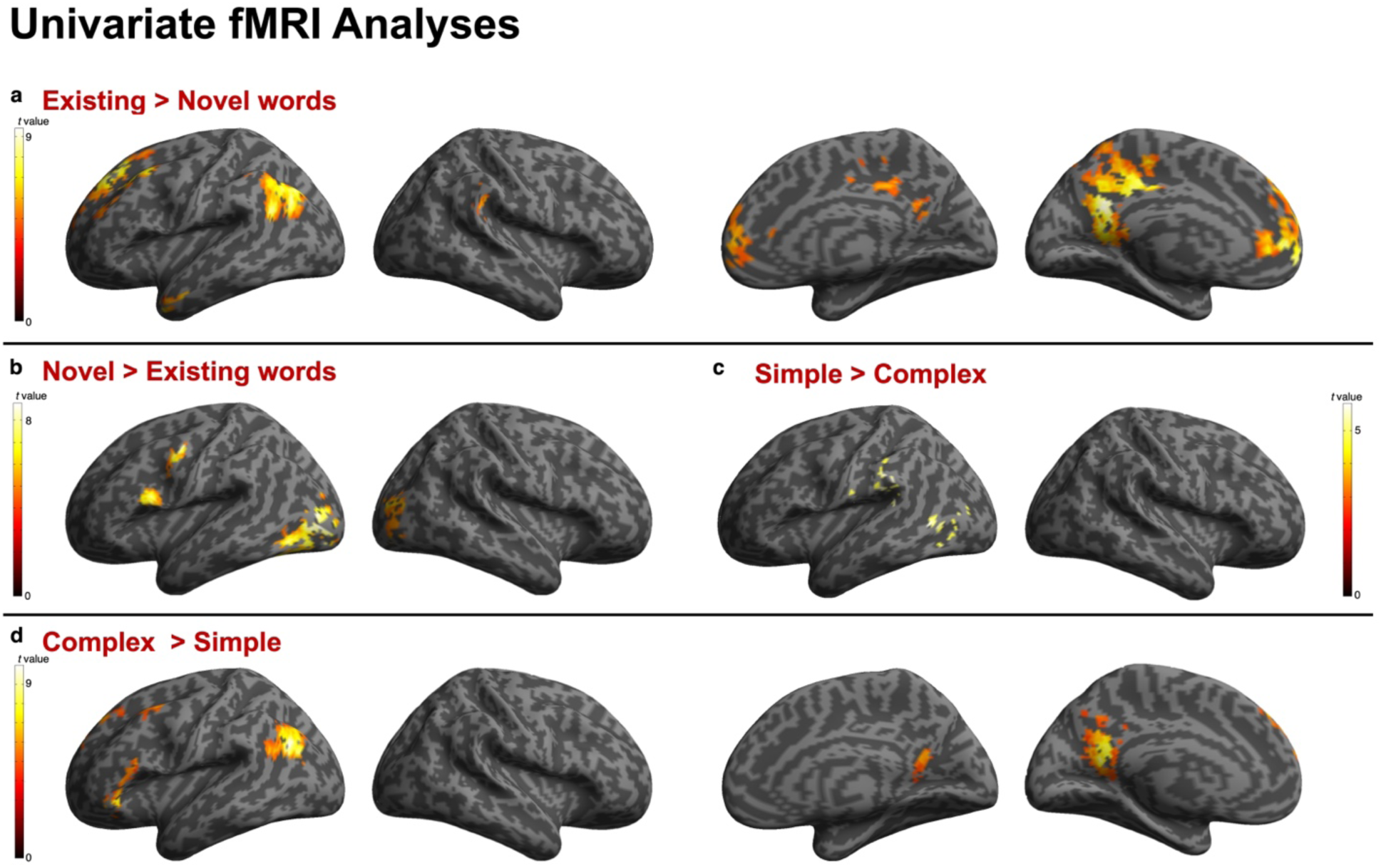
| Simple effects in univariate fMRI data. Contrast estimates and renderings of the simple effects assessing the Lexicality (a. and b.) and Morphological complexity (c. and d.) factors in univariate fMRI data. Colorbars indicate *t* values.

Significant interaction effects (**Fig. 4, Supplementary Table S9**) were detected in brain areas that–with the only exception of the cerebellum–are part of the semantic network^16^ (**Supplementary Table S2**), suggesting continuities in the underpinnings of semantic activation for existing words and novel words that embed morphemes. More specifically, we observed greater activation for existing words (regardless of their morphological structure) and morphologically complex novel words compared to morphologically simple novel words in the triangular portion of the left inferior frontal gyrus, in the left medial superior frontal gyrus, in the left middle temporal gyrus, in the left angular gyrus, and in the right cerebellum.

**Fig. 4.**
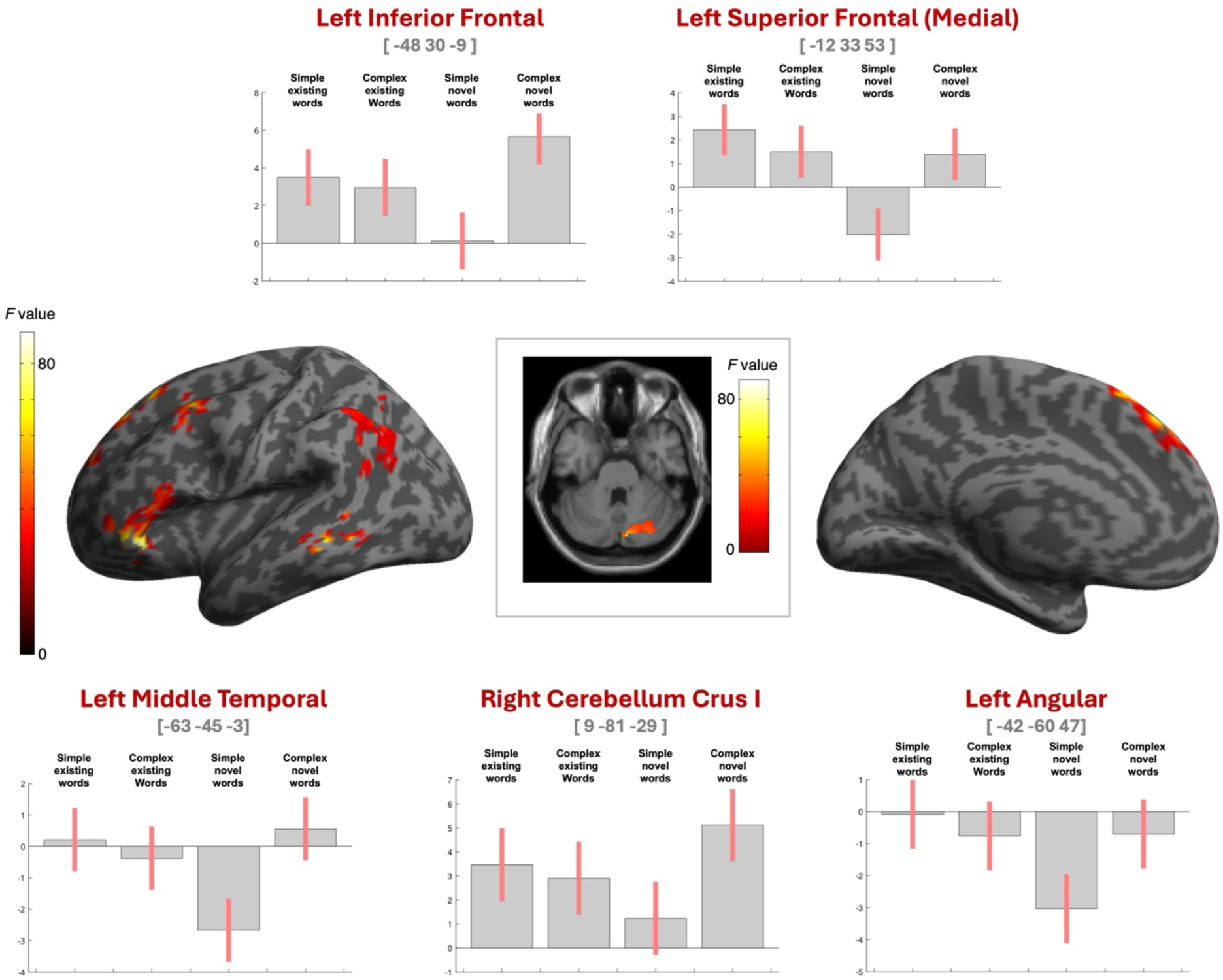
| Lexicality by Morphological complexity interaction in univariate fMRI data. Contrast estimates (bar plots) and renderings of the Lexicality-by-Morphological complexity interaction in univariate fMRI data. Number triplets below anatomical labels represent MNI coordinates (x,y,z) in mm for the activation peak within each cluster. Colorbars indicate *F* values. Error bars indicate 90% confidence intervals.

### Results of the Representational Similarity Analysis

We adopted the FastText^54^ model to induce high-dimensional computational estimates of semantic representations for existing and novel words (**Fig.1d)**. FastText builds on patterns of statistical co-occurrences in a large textual corpus (in this case Common Crawl and Wikipedia) to induce semantic representations for existing and novel words via their constituting *ngrams*^54, 61, 62^. In line with distributional semantic models^74^, FastText operationalizes meaning as a set of coordinates in a high-dimensional space, in which each dimension (300 in our case) provides a proxy measure of the relevance of the word in a (abstracted) linguistic context.

To explore the extent to which the neural underpinnings of semantic access for existing and novel words align with such high-dimensional representations, we performed a RSA (**Fig.1g**). More specifically, we conducted RSA within an orthogonal semantic mask (see Methods for details). This was done to locate the areas in the brain whose pattern of activity during stimuli presentation show isomorphisms with patterns of semantic activation operationalized by the computational semantic model, and to test whether the extent of such isomorphisms show differences between stimuli. Notably, this fine-grained semantic analysis was carried out after partialling-out orthographic similarity between stimuli (as measured through Levenshtein Distance), while comparing stimuli categories by adopting the same 2-by-2 (Lexicality by Morphological complexity) design adopted for univariate analyses. Results revealed that, compared to novel words, patterns of brain activation for existing words better aligned with the computational semantic model in a set of areas encompassing the left precuneus, left middle temporal gyrus, and left inferior parietal gyrus (**Fig.5a**, **Supplementary Table S10**), with fine-grained sematic representations for morphologically simple existing words mapped in the left precuneus (**Figure 5b**, **Supplementary Table S11**). The patterns of activation of a set of areas involving superior and middle frontal cortices as well as the precuneus showed greater alignment with the computational semantic model for morphologically complex stimuli than for morphologically simple stimuli (**Figure 5c**, **Supplementary Table S12**), with representations for complex existing words—mapping in the left middle and superior frontal gyri (**Figure 5d**, **Supplementary Table S13**)—showing greater alignment with the model than morphologically simple existing words (**Figure 5e**, **Supplementary Table S14**). Critically, brain-semantic model representational similarity was found to be greater for morphologically complex novel words than simple novel words in areas mainly involving the left precuneus (**Figure 5f**, **Supplementary Table S15**).

**Fig. 5.**
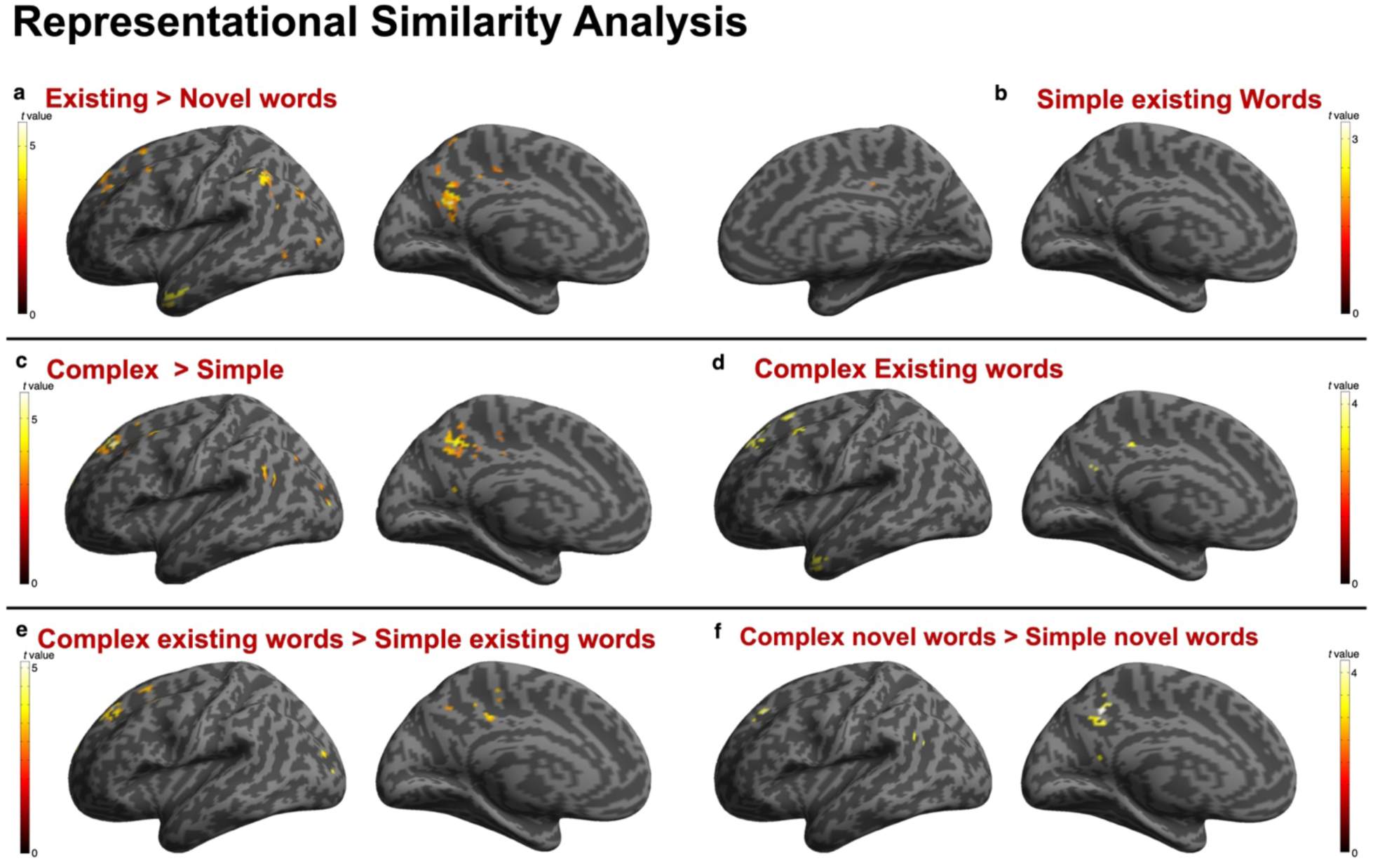
| Representation Similarity Analysis results. Contrast estimates and renderings of significant effects in the Representation Similarity Analysis. Colorbars indicate *t* values.

Regression analyses revealed a significant effect of orthographic neighborhood (*t*(188)=-4.180, *p*< 0.001), suggesting lower activity in these areas for stimuli with a denser orthographic neighborhood, which further confirms the semantic (as opposed to orthographic) nature of the brain areas under consideration.

## Discussion

Through a combination of behavioral, computational, and univariate and multivariate neuroimaging techniques, we explored whether (and to what extent) novel words activate the neurocognitive hallmarks of semantic processing typically involved in reading existing words. As anticipated by a considerable wealth of literature, we did detect significant lexicality effects in both behavioral and fMRI data^11, 12, 71, 75–78^. These effects point to the fact that existing and novel words (in principle considered as two distinct categories of stimuli) trigger different neurocognitive underpinnings: existing words entailed greater involvement of areas typically associated with lexical-semantic mechanisms (as indicated by greater activation than novel words in the left angular gyrus, precuneus, superior frontal gyrus, middle temporal gyrus, and supramarginal gyrus^11, 12, 24^), whereas novel words more greatly engaged early visuo-orthographic (as indicated by the involvement of middle occipital cortices bilaterally^14, 28, 29, 79–81^) and phonological (as indicated by the involvement of the inferior frontal gyrus^24, 82, 83^) processing. *Prima facie*, these results seem to suggest that existing and novel words are qualitatively different in the neurocognitive mechanisms they engage. However, the primary focus of this study was to explore the possibility that – underneath the “existing word” vs “novel word” dichotomization – there could be a set of partially overlapping mechanisms pointing to semantics, as the very core that any linguistic stimulus, irrespective of its lexical status, will prompt. Our results support this possibility.

Both the behavioral metric and univariate fMRI data activation was maximal for existing words (regardless of their morphological complexity), comparatively lower (more specifically, similar in fMRI data, lower in behavioral Overall Semantic Loading data) for morphologically complex novel words, and minimal for morphologically simple novel words. The topography of this effect in brain data points towards anatomical structures involved in semantic processing: the left angular gyrus (whose activity has been associated with information integration and knowledge retrieval^16, 84, 85^) the left middle temporal gyrus (whose impairment has been related to language comprehension and semantic deficits^86–90^), the left medial superior frontal gyrus (previously associated with goal-directed retrieval of semantic information^16, 91^), the left inferior frontal gyrus (ubiquitously activated in language tasks, important for phonology and processing efficiency^16, 75, 82, 92–95^) and the right cerebellum (whose involvement in semantics has been related to predictive processes^67–70^).

Beyond the average degree of activation of the semantic system, the RSA offers insight on the organization of the activation patterns within the semantic system. Data indicate that letter strings that can rely on familiar meaning-bearing units (i.e., words, morphemes) capitalize on fine-grained representations (that we gauged through the FastText^54^ computational semantic model) which can be mapped in high-order areas involved in the processing and integration of meaning. The representational advantage of existing words over novel words was significant in a default-mode network area critically involved in imagery such as the left precuneus^96–99^, in the left temporal pole, which is regarded as a primary semantic hub in the brain^17, 100, 101^, and in the left inferior frontal gyrus. The advantage in fine-grained semantic encoding of morphologically complex stimuli over simple ones was localized in left frontal cortices, whereas the advantage of morphologically complex novel words over morphologically simple novel words was significant in the left precuneus.

These data are particularly interesting to relate to previous theoretical standpoints of lexical and semantic processing. Localist models of the human lexical-semantic system have formalized the process of meaning extraction from print while assuming the existence of a mental representation for each known word. Although differences exist between serial^19–22^ and cascaded^23^ accounts on how activation spreads between the lexical and the semantic level of stimuli processing, these frameworks agree in that novel words are essentially meaningless, as there is no “memory” of their meaning to be retrieved. The present findings contradict this position: participants in our study were able to activate meaning so as to provide valence and imageability judgements even for words they had never encountered before. Critically, our results indicate that participants activated meaning by tapping into neurocognitive mechanisms comparable to those engaged in semantic processing for words.

Our results dovetail hints from literature suggesting some degree of continuity in semantic processing for existing and novel words^58–61, 63, 102^. Our findings push this suggestion to the level of description that includes the interface between cognitive and neural processes. In particular, our data indicate that semantic activation for novel words can be partially overlapping to that of existing words, provided that novel words contain morphemes that can point to meaning. These results suggest a set of functional neurocognitive mechanisms for semantic processing that overlap (at least partially) between existing and novel words. Accordingly, the linguistically-normative status of “existing” word vs. “novel” word does not seem to be the primary determinant of semantic activation: sizeable activation of the semantic system can be detected whenever the incoming input contains a relatively salient unit of form that reliably points to meaning. Such units of form could take the shape of words or morphemes, and their availability in an incoming stimulus allows reliance on fine-grained semantic representations.

This scenario fits well with the idea that there is no “brick wall” that separates existing words from novel words in the human mind and brain, but rather a gradient, a gentle hill that spans from one linguistic type of stimulus to the other. Such gradient fits well with distributed conceptualizations of linguistic knowledge that bypass the need of positing the existence of a “lexicon”^34, 44, 46, 50^, in that the boundary between two representations is quantitative rather than qualitative. In this landscape, morphemes play a pivotal role, in that they constitute a gateway to meaning even when words that contain them are unattested. This complements previous literature on morphological processing in suggesting that, although morphological decomposition is ultimately due to sensitivity to superficial orthographic mechanisms^103^, the nature of the information they convey is inherently semantic^63^.

The pivotal role of morphemes as interface between form and meaning aligns with the idea that language is less arbitrary than traditionally thought^104^. Contrary to the classical proposal that there is no apparent link between form and meaning^105^ (i.e., linguistic arbitrariness), it was pointed out that there are reliable statistical associations between sublexical elements and semantic features (i.e., “systematicity”^106^), as well as relations that consistently bind certain linguistic sounds to meaning (i.e., “sound-symbolism”, or “iconicity”^107–109^). In this context, morphemes represent a specific case of systematicity: they are elements that stand out by virtue of their statistical properties and drive semantic access^65, 66^. Our data indicating reliance on morphology as a gateway to meaning while processing novel words mirror evidence indicating that statistically relevant patterns are extensively relied upon in novel word learning^110–113^. Strengthened by the fact that humans are able to locate statistical regularities since early in development^114^, this result suggests that cues that systematically associate form and meaning represent the most convenient way to “make sense” of a novel word.

In summary, the present results indicate that, even if a letter string is not (yet) a word “to us” it can still be a word “in us” — in terms of recruitment of similar neurocognitive substrates. In this landscape, the boundaries between existing words and novel words are more nuanced than believed until nowadays, as the possibility for written stimuli to trigger semantic processing ultimately depends on whether they contain form-related information that reliably points to meaning, rather than on their lexical status.

The present study invites a reconsideration of the extent to which words should be considered a psychological entity, in that the capability for stimuli to activate mental representations appears to be independent from lexicality, in formal sense. Still, our results reconcile previous literature describing lexical effects in language processing^11, 12, 71, 75–78^ with the everyday ecological observation that we are indeed able to understand (and learn^1^) “meaningable” words, even when they are completely unfamiliar to us. As humans, we cannot indeed refrain from making sense of the stimuli we come across in the outside world^115^, regardless of whether we have encountered them before or not. Novel words—and the meaning they convey—represent an ideal case study to explore such defining property of our species.

## Methods

### Participants

A total of 32 volunteers (20 females) took part to the study (*mean* age = 24.753 years, *SD* = 3.952), recruited through the local University participant recruitment platform, flyers, and word of mouth. Participants reported no vision, neurological, or psychiatric disorder. All participants were right-handed according to the Edinburgh Handedness Inventory^116^ (*mean score* = 0.941, *SD* = 0.09). Participants could choose to be compensated for their participation either with a with a 15€ reimbursement, or with course credits. The study was carried out in accordance with the Helsinki declaration. The study was approved by the Ethical Committee of the University of Milano-Bicocca (approval code: 589-2021).

### Stimuli

We devised a set of stimuli to be administered to Italian speakers, comprising morphologically complex existing words, morphologically simple existing words, morphologically complex novel words and morphologically simple novel words. The initial set consisted of 60 stimuli per type. More specifically, morphologically complex existing words were selected from the training set used by Bonandrini et al.^63^ (in turn extracted from the DerIvaTario corpus^117^) for computational modelling of affixation. After dropping words for which affixation entailed an orthographic change in the stem (i.e., *tassista* [stem= *taxi*], *turista* [stem= *tour*], *umorismo* [stem= *humor*], *cavaliere* [stem= *cavallo*], *marinaio* [stem= *mare*]) and redundant items, we selected the 6 most productive non-evaluative affixes that entail a noun-to-noun transformation (e.g., *arte*(= art, noun) + (*-ista*) = *artista* (= artist, noun)): –*ista*, –*ismo*, –*iere*, –*iere*, –*eria*, –*aio*). For each affix, we selected the 10 most frequent affixed words from the set. As far as morphologically complex novel words are concerned, we selected stems from the Subtlex-it^118^ corpus while avoiding loan words, proper and geographical names, taboo words and words related to religion, and plural items, so that their length and frequency distribution (as well as the length of the affixed form) matched that of the subset of items with each affix (all *p*s ≥ 0.560). Morphologically simple word stimuli were extracted from the set by Vergallito et al.^119^. We discarded items including accents and kept only morphologically simple nouns whose length (*W*= 1660*, p*= 0.441) and frequency (*W*= 2040, *p=* 0.209) matched that of morphologically complex words. Morphologically simple novel words were also taken from the Wuggy-generated^120^ set by Vergallito et al.^119^. More specifically, we dropped items ending with derivational or inflectional morphemes and extracted items whose length matched that of affixed novel words (*W=*2100, *p*= 0.101).

A second selection step was carried out on this set of stimuli through a pilot study conducted on 24 volunteers (6 males, mean age= 23.375 years, sd= 3.910). In particular, for each category of stimuli, the 12 items that yielded the greatest inter-individual uncertainty (i.e., standard deviation) in behavioural ratings (i.e., imageability and valence) with reference to the centre of the bi-dimensional response grid were discarded. The final set of stimuli (48 per category) was matched across categories according to length, in that there was no significant effect of Lexicality (*F*(1,188)= 1.257, *p*= 0.264), Morphological complexity (*F*(1,188)= 1.570, *p*= 0.212) or an interaction effect (*F*(1,188)= 0.213, *p=*0.645). Morphologically simple and complex existing words did not statistically differ according to frequency (*W*= 897.5, *p*= 0.063). Morphologically complex existing and novel words did not differ according to stem frequency (*W*= 1179*, p*= 0.846). Within morphologically complex stimuli, the distribution of affix use did not differ between existing and novel words (*χ*^2^(5)= 0.877, *p*= 0.972).

### Task and procedure

After signing informed consent, and a familiarization phase with the task outside the MRI scanner, experimental stimuli were administered across 6 experimental runs, each lasting approximately 9 minutes. During each run, 32 stimuli were presented in blocks containing existing or novel words. Before each block, participants saw a message (for 9.7 seconds) indicating whether the forthcoming stimuli would be existing or novel words. During each trial, a fixation cross was presented for a jittered inter-stimulus interval (ISI) ranging from 2.5s to 7.5s following the proportions suggested by Dale et al.^121^, after which the stimulus was presented (in uppercase font) for 2.8s. Another fixation cross was subsequently presented for a jittered ISI between 1.5s and 2.5s with the same proportions as above. This was followed by the imageability and valence grid, which remained on screen for a fixed time of 8s, during which participants were asked to provide their ratings. To do so, participants operated on two MRI-compatible NordicNeurolab ResponseGrip button boxes (one per each hand, each one with a button for the thumb and one for the index finger). The right-hand buttons were used to move the cursor along the imageability axis, while the left-hand buttons were used to move the cursor along the valence axis. Visual stimuli were presented by means of a 40” MRI-compatible NordicNeuroLab InRoom Viewing System located behind the MRI bore and a mirror mounted on the MRI head coil. The experimental paradigm was created and administered through PsychoPy (v. 2022.2.4)^122^.

### MRI data acquisition

MRI scans were acquired with a 3 Tesla Philips Ingenia 3.0T CX MRI scanner using a 32-channels head coil. During the experimental session, whole-brain functional images were acquired with a T2*-weighted gradient-echo, EPI pulse sequence, using BOLD contrast (TR = 2000 ms, TE = 30 ms, flip angle = 75°, SENSE factor = 2). Each functional image comprised 35 axial slices (3 mm isotropic voxel size; FOV = 240 x 240 mm, inter-slice spacing= 0.3 mm). Each participant underwent 6 consecutive functional scanning sessions, each comprising 274 scans, for a total duration of about 9 min each. At the end of each experimental session, we obtained a 3D T1-weighted turbo field echo anatomical scan (243 sagittal slices, TR = 12 ms, TE = 5 ms, flip angle = 8°; FOV = 269 × 269 mm, 0.7 mm isotropic voxel size).

### Data analysis

<u>Behavioral data:</u> We initially conducted a sanity check to make sure that participants’ responses were not converging towards the center of the axes of the two-dimensional grid. To do so, separately for each participant, we computed the Euclidean norm of the imageability and valence judgements for each stimulus *i* as follows:

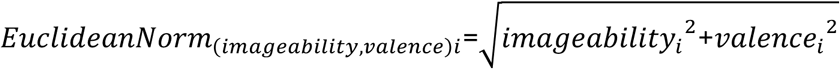

We then ran four linear mixed-effects models^123, 124^ (one per each category of stimulus) on the Euclidean Norm to check that participants’ responses were not anchored to the origin of the axes. More specifically, we evaluated the significance of the fixed-effects intercept in the four models.

The effects of Morphological complexity, Lexicality and their interactions were then estimated on valence and imageability judgements by means of linear mixed-effects models.

To obtain a global behavioral measure of semantic activation, we computed a metric that we defined “Overall Semantic Loading”, which captures the Euclidean Distance from the “semantic zero” point in the two-dimensional semantic space adopted to probe participants’ intuitions on stimuli meaning. The “semantic zero” point represents the absence of semantic activation, and in the context of our task it corresponds to minimal imageability (i.e., –3 in the rating grid) and null valence (i.e., 0 in the rating grid) in our two-dimensional semantic space. Separately for each participant, the OSL metric for each stimulus *i* was computed as follows:

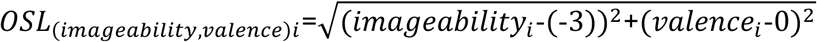

Hence:

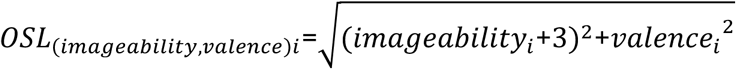

The effects of Morphological complexity, Lexicality and their interactions were estimated on OSL by means of a linear mixed-effects model. Departure from the “semantic zero” point was evaluated through the significance of the fixed-effects intercept in four linear mixed-effects models (one per each category of stimulus). All linear mixed-effects models were run in R (version 4.3.3) via the lme4 package^123^, and contained (in addition to the fixed effects described above) by-participants and by-stimulus random intercepts.

<u>fMRI preprocessing:</u> After conversion from the DICOM to the NIfTI format using dcm2niix (https://www.nitrc.org/plugins/mwiki/index.php/dcm2nii:MainPage), all structural and functional volumes from each participant were visually inspected using FSLEyes (https://open.win.ox.ac.uk/pages/fsl/fsleyes/fsleyes/userdoc/). Subsequently, volumes were manually reoriented using SPM12 (v. 7771)^125^ (https://www.fil.ion.ucl.ac.uk/spm/software/spm12/) in MATLAB (v.2023b https://www.mathworks.com/products/matlab.html). Scan-to-scan signal variance was visually inspected using TSDiffAna (https://www.fil.ion.ucl.ac.uk/spm/ext/#TSDiffAna). Temporal signal-to-noise ratio measures were acquired through MRIQC^126^ (https://mriqc.readthedocs.io/en/latest/). Preprocessing of structural and functional data was conducted with fMRIPrep (v.23.1.0)^127^ (https://fmriprep.org/en/stable/), which is based on Nipype 1.8.6^128^. The anatomical T1-weighted image was corrected for intensity non-uniformity with N4BiasFieldCorrection^129^ (distributed with ANTs^130^), and used as T1w-reference throughout the workflow. The T1w-reference was skull-stripped with a Nipype implementation of antsBrainExtraction.sh (from ANTs), using OASIS30ANTs as target template. Brain tissue segmentation of cerebrospinal fluid (CSF), white-matter (WM) and gray-matter (GM) was performed on the brain-extracted T1w using fast^131^ in FSL. Brain surfaces were reconstructed using recon-all (FreeSurfer 7.3.2^132^), and the previously estimated brain mask was refined with a custom variation of the method to reconcile ANTs-derived and FreeSurfer-derived segmentations of the cortical gray-matter of Mindboggle^133^. Volume-based spatial normalization to two the MNI152 standard space was performed through nonlinear registration with antsRegistration, using a brain-extracted version of the T1w template.

As for the preprocessing of functional data, for each of the 6 BOLD runs per subject, the following preprocessing was performed. First, a reference volume and its skull-stripped version were generated using a custom methodology of fMRIPrep. Head-motion parameters with respect to the BOLD reference (transformation matrices, and six corresponding rotation and translation parameters) are estimated before any spatiotemporal filtering using mcflirt^134^ in FSL. The BOLD time-series (including slice-timing correction when applied) were resampled onto their original, native space by applying the transforms to correct for head-motion. These resampled BOLD time-series will be referred to as preprocessed BOLD in original space, or just preprocessed BOLD. The BOLD reference was then co-registered to the T1w reference using bbregister (FreeSurfer) which implements boundary-based registration^135^. Co-registration was configured with twelve degrees of freedom, to account for distortions remaining in the BOLD reference. Several confounding time-series were calculated based on the preprocessed BOLD: framewise displacement (FD), DVARS and three region-wise global signals. FD was computed using two formulations following Power^136^ (absolute sum of relative motions) and Jenkinson^134^ (relative root mean square displacement between affines). FD and DVARS are calculated for each functional run, both using their implementations in Nipype (following the definitions by Power et al.^136^). The three global signals are extracted within the CSF, the WM, and the whole-brain masks. Additionally, a set of physiological regressors were extracted to allow for component-based noise correction (CompCor)^137^. Principal components are estimated after high-pass filtering the preprocessed BOLD time-series (using a discrete cosine filter with 128s cut-off) for the two CompCor variants: temporal (tCompCor) and anatomical (aCompCor). tCompCor components are then calculated from the top 2% variable voxels within the brain mask. For aCompCor, three probabilistic masks (CSF, WM and combined CSF+WM) are generated in anatomical space. The implementation differs from that of Behzadi et al.^137^ in that instead of eroding the masks by 2 pixels on BOLD space, a mask of pixels that likely contain a volume fraction of GM is subtracted from the aCompCor masks. This mask is obtained by dilating a GM mask extracted from the FreeSurfer’s aseg segmentation, and it ensures components are not extracted from voxels containing a minimal fraction of GM. Finally, these masks are resampled into BOLD space and binarized by thresholding at 0.99 (as in the original implementation). Components are also calculated separately within the WM and CSF masks. For each CompCor decomposition, the k components with the largest singular values are retained, such that the retained components’ time series are sufficient to explain 50 percent of variance across the nuisance mask (CSF, WM, combined, or temporal). The remaining components are dropped from consideration. The head-motion estimates calculated in the correction step were also placed within the corresponding confounds file. The confound time series derived from head motion estimates and global signals were expanded with the inclusion of temporal derivatives and quadratic terms for each^138^. Frames that exceeded a threshold of 0.5 mm FD or 1.5 standardized DVARS were annotated as motion outliers. Additional nuisance timeseries are calculated by means of principal components analysis of the signal found within a thin band (crown) of voxels around the edge of the brain^139^. The BOLD time-series were resampled into standard space, generating a preprocessed BOLD run in the MNI152 space. First, a reference volume and its skull-stripped version were generated using a custom methodology of fMRIPrep. All resamplings can be performed with a single interpolation step by composing all the pertinent transformations (i.e. head-motion transform matrices, susceptibility distortion correction when available, and co-registrations to anatomical and output spaces). Gridded (volumetric) resamplings were performed using antsApplyTransforms (ANTs), configured with Lanczos^140^ interpolation to minimize the smoothing effects of other kernels. Non-gridded (surface) resamplings were performed using mri_vol2surf (FreeSurfer). Many internal operations of fMRIPrep use Nilearn 0.10.1^141^, mostly within the functional processing workflow. Volume spatial smoothing was eventually applied in SPM12 (FWHM 6×6×6 mm).

<u>Univariate fMRI Analyses:</u> Smoothed functional volumes from each participant were included in analyses at the individual participant level. More specifically, we specified a general linear model (comprising six separate runs) including regressors of interest for each of the four categories of stimuli (i.e., combination of the factors “Lexicality” and “Morphological complexity”), the evaluation grid (involving the time interval in which participants produced the motor responses for imageability and valence ratings), and task instructions events. The model included also covariates of no interest (i.e., the 6 realignment parameters) representing nuisance variables derived from the pre-processing. The “Global normalization” parameter in SPM12 was set to “None”. Participant-level contrasts consisted of simple effects estimated for each of the four stimuli categories of interest (i.e., weight=1 for each condition of interest, 0 otherwise), which were then included in a group-level factorial analysis to explore whole-brain effects of Lexicality, Morphological complexity and their interaction. A peak-level threshold of p< 0.05 Family-Wise Error (FWE) was adopted, and a cluster extent threshold of at least 30 voxels.

Overlap of the effects of Lexicality, Morphological complexity (as well as their interaction) with the putative location of the semantic system in the brain was evaluated as follows: the functional Activation Likelihood Estimation meta-analytic map representing “all semantic foci” in Binder et al.^16^ was transformed from the Talairach and Tournoux^142^ to the Montreal Neurological Institute (MNI152) atlas space, by first warping a T1-weighted structural image from the former to the latter space using the CAT12^143^ segmentation algorithm (version 12.9), and by then applying the obtained transformation matrix to the functional meta-analytic map. Then, for each significant cluster within each effect, the number of voxels (and their percentage, with reference to each cluster’s total number of voxels) overlapping with the reference semantic map by Binder et al. was computed.

<u>RSA:</u> Un-smoothed functional volumes from each participant were included in participant-level analyses including one regressor of interest for each stimulus and nuisance variables^144^. Contrasts were set so as to produce a Tmap for each stimulus for each participant. Stimuli-wise Tmaps were used as input for RSA in CosmoMVPA^145^ (v.1.1.0) in MATLAB 2023b, which were conducted (separately for each of the four categories of stimuli) by using the matrix of semantic (cosine)^146^ similarity of stimuli as computed from FastText^54^ (pre-trained in Italian on Common Crawl and Wikipedia, using the “continuous bag of words” algorithm with position-weights, 300 dimensions, with character n-grams of length 5, a window of size 5 and 10 negatives; https://fasttext.cc/docs/en/crawl-vectors.html) embeddings as a model matrix, after partialling-out orthographic similarity between stimuli, as measured through Levenshtein Distance. FastText was chosen as reference semantic model due to its capability to gauge semantic effects for both existing and novel words^54, 55, 59, 63^.

The RSA was conducted with a searchlight approach (with a neighborhood with constant number of voxels= 30 voxels and variable radius to minimize estimation biases due to differences in regional anatomy) within a mask containing voxels resulting from either the “existing word> novel word” or the “novel word> existing word” contrasts in univariate analyses (each one thresholded at p< 0.05 FWE-corrected peak-level and k>30). Spearman correlation was adopted to compute the RSA. After Fisher transformation, RSA data for each category of stimulus for each participant was entered in a group-level full-factorial analysis in SPM (in line with univariate fMRI analyses, see previous paragraph). A peak-level threshold of p< 0.05 Family-Wise Error (FWE) was adopted. For additional exploratory purposes, we also inspected results with peak-level p< 0.001 uncorrected threshold.

## Acknowledgements

M.M. and R.B. were supported by the European Union (ERC-COG-2022, BraveNewWord, 101087053). The views and opinions expressed are those of the authors alone and do not necessarily reflect those of the European Union or the European Research Council Executive Agency. Neither the European Union nor the granting authority can be held responsible for them.

## Author contributions

The authors made the following contributions: R.B: Methodology, Formal analysis, Investigation, Data Curation, Writing – Original Draft, Visualization; S.A: Conceptualization, Methodology, Resources, Writing – Review & Editing; S.S: Conceptualization, Methodology, Resources, Writing – Review & Editing; G.B: Investigation, Writing – Review & Editing; M.M: Conceptualization, Methodology, Writing – Review & Editing, Supervision; M.T.: Conceptualization, Methodology, Formal analysis, Investigation, Data Curation, Writing – Review & Editing, Supervision.

## Competing interests

The authors declare no competing interests.

## Supplementary Materials

**Supplementary Table S1.**
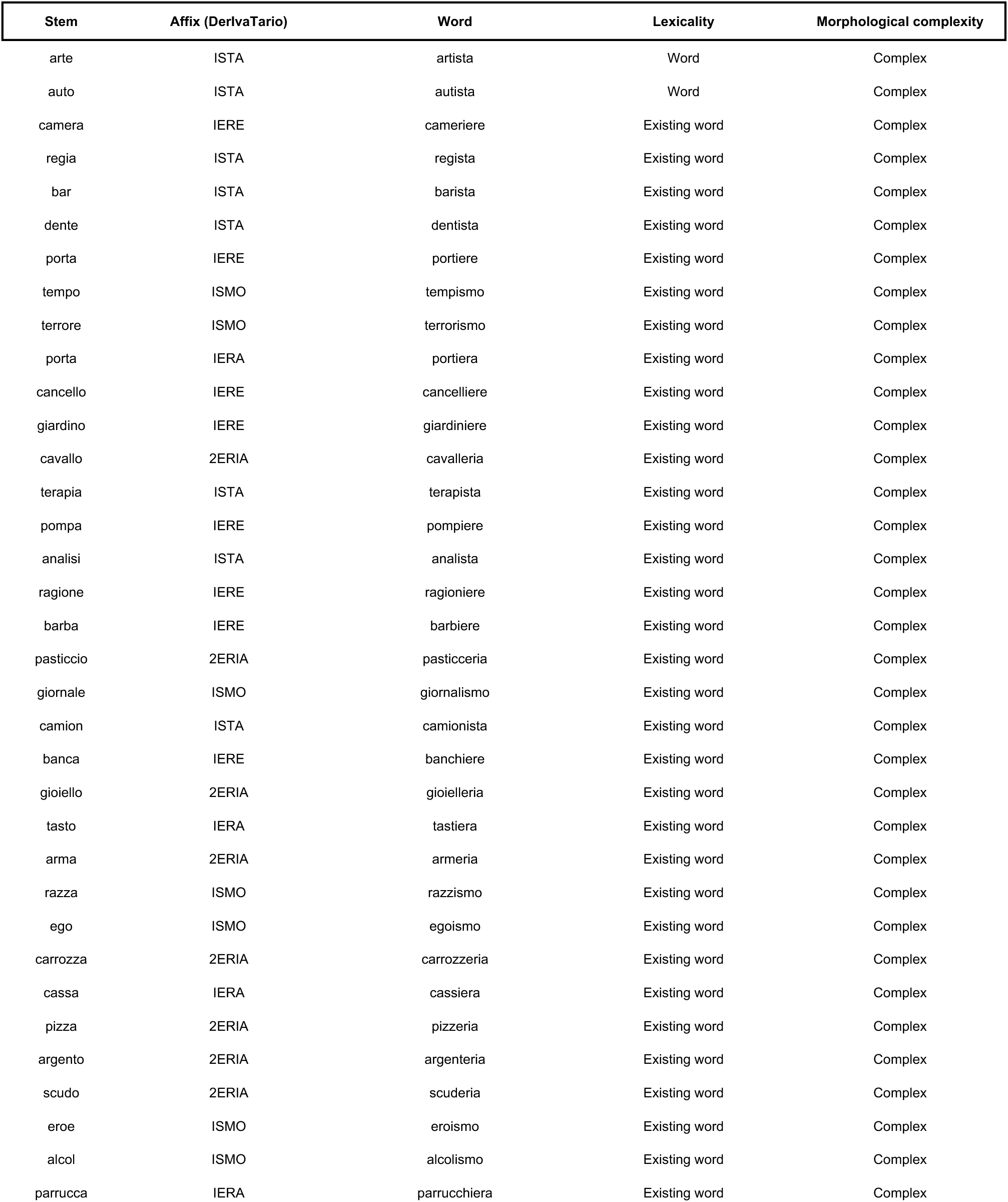
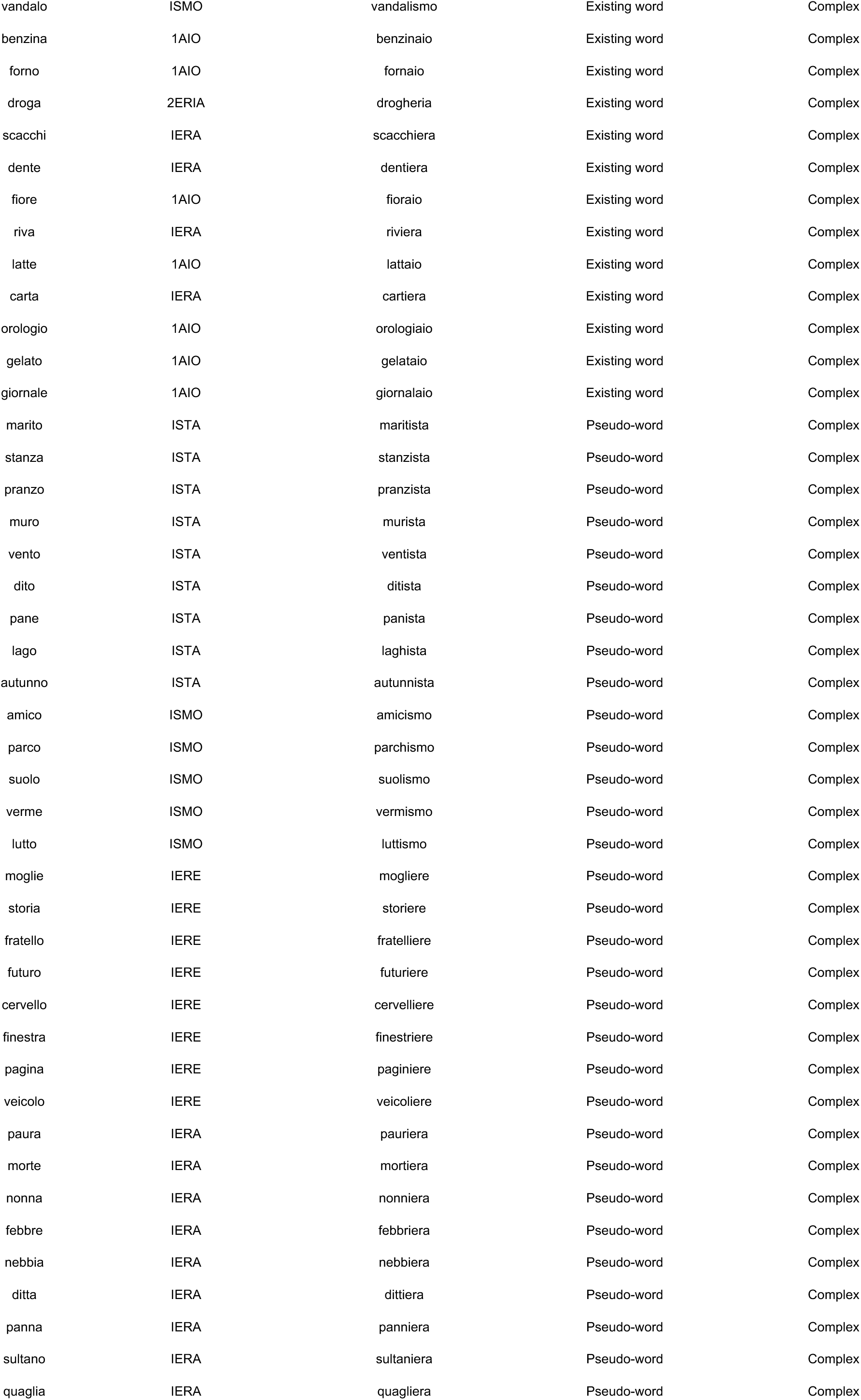
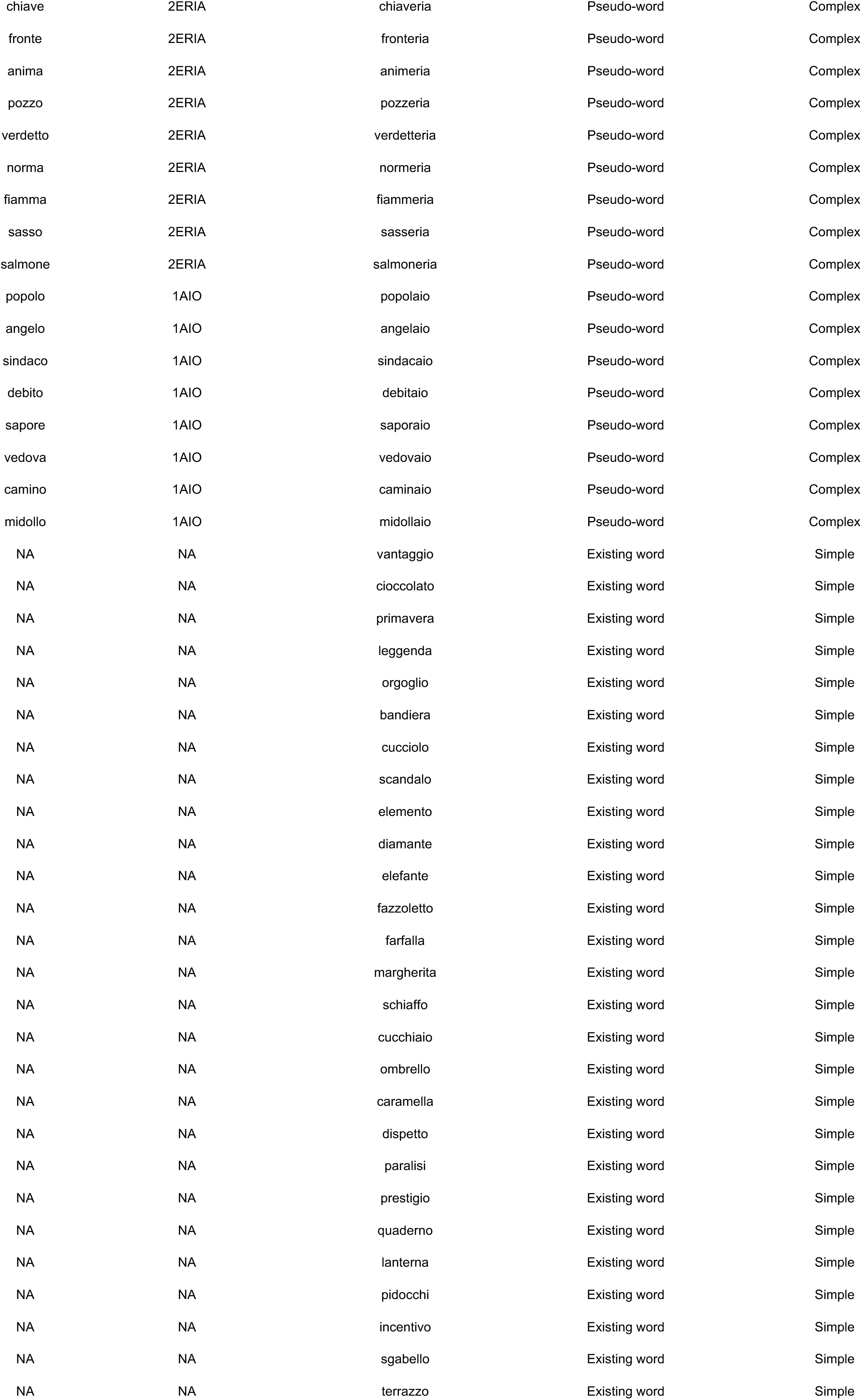
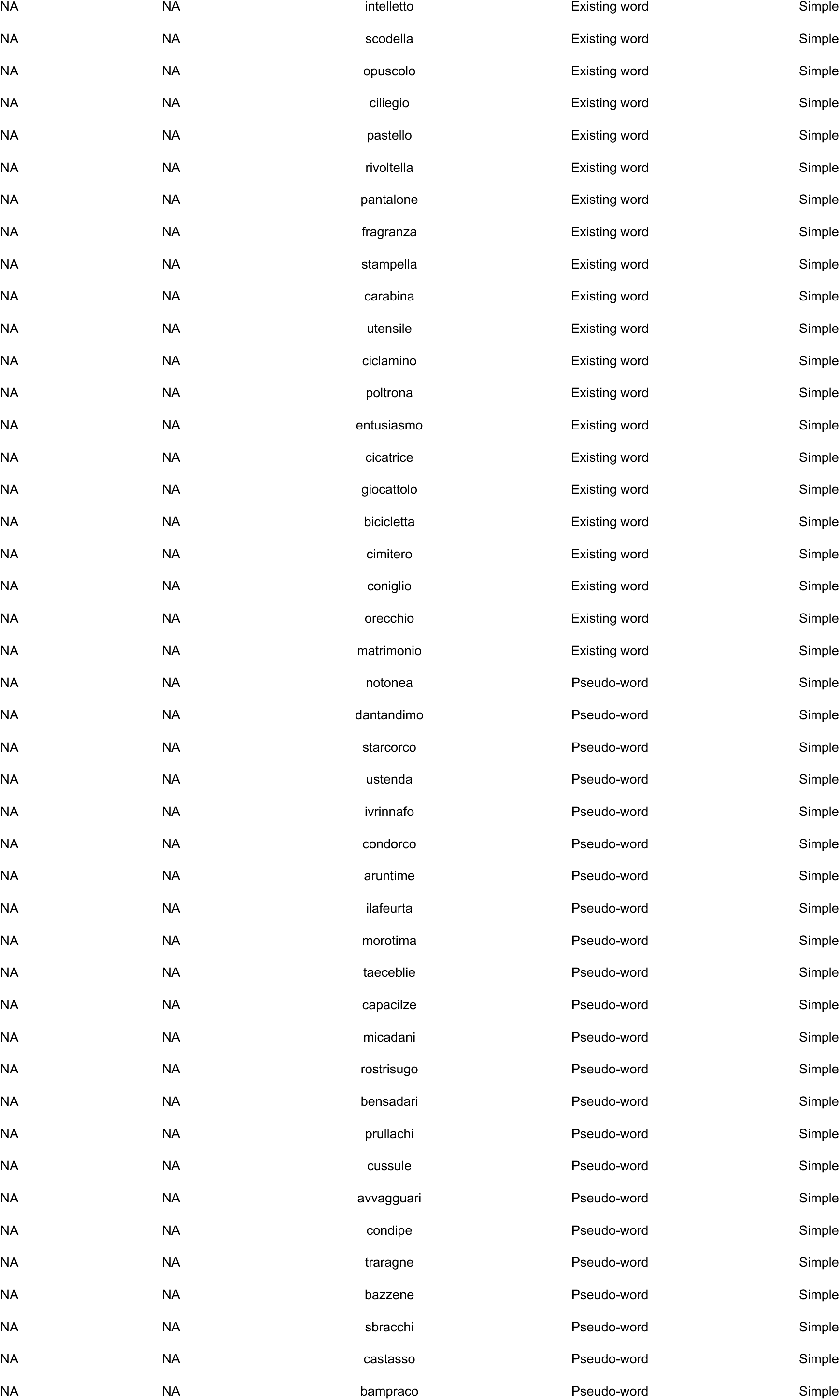
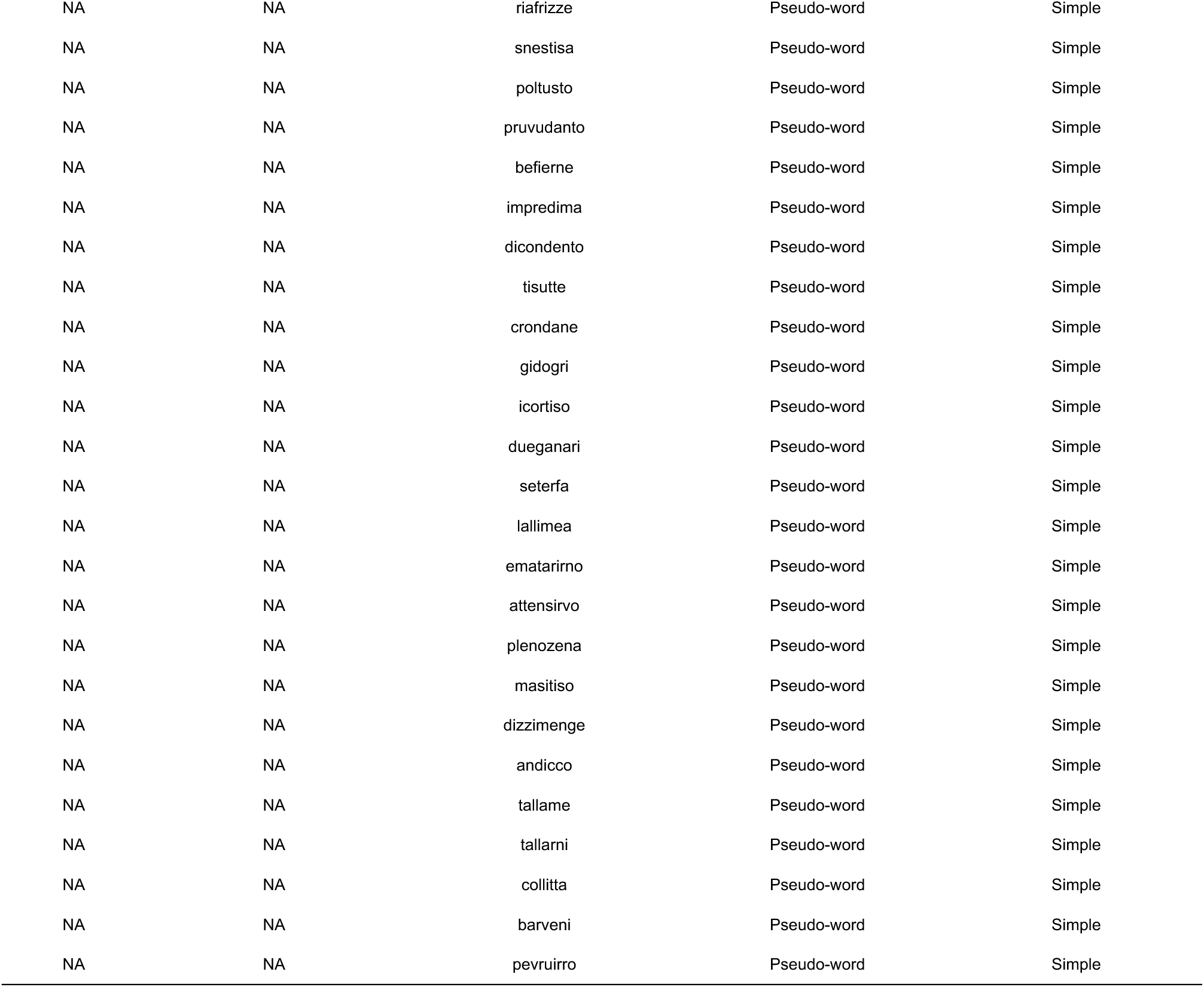
| Overview of experimental stimuli. Please note that affixes are reported as in the DerIvaTario^1^ corpus. Numbers before affixes denote the existence of alternative versions of the same affix.

**Supplementary Table S2.** | Overlap with the semantic system. Overlap between significant effects in univariate fMRI analyses and the semantic system as characterized by Binder et al.^2^ The column “k” indicates the total number of voxels in that cluster. The column “% overlapping voxels” indicates, out of the total number of voxels in each cluster the percentage of voxels overlapping with the reference semantic meta-analytical map.

| Effect | Cluster | Location | k | # overlapping voxels | % overlapping voxels |
| --- | --- | --- | --- | --- | --- |
| Lexicality | 1 | Left superior frontal gyrus | 896 | 218 | 24.33 |
| Lexicality | 2 | Left precuneus | 641 | 203 | 31.669 |
| Lexicality | 3 | Left angular gyrus | 319 | 238 | 74.608 |
| Lexicality | 4 | Left middle occipital gyrus | 284 | 0 | 0 |
| Lexicality | 5 | Left inferior frontal gyrus (pars opercularis) | 133 | 1 | 0.752 |
| Lexicality | 6 | Right middle occipital gyrus | 80 | 0 | 0 |
| Lexicality | 7 | Left middle temporal gyrus | 33 | 6 | 18.182 |
| Morphology | 1 | Left precuneus | 227 | 154 | 67.841 |
| Morphology | 2 | Left angular gyrus | 224 | 181 | 80.804 |
| Morphology | 3 | Left superior frontal gyrus | 196 | 73 | 37.245 |
| Morphology | 4 | Left inferior frontal gyrus (pars triangularis) | 105 | 58 | 55.238 |
| Morphology | 5 | Left supramarginal gyrus | 54 | 3 | 5.556 |
| Lexicality by Morphology | 1 | Left medial superior frontal gyrus | 396 | 93 | 23.485 |
| Lexicality by Morphology | 2 | Left inferior frontal gyrus (pars triangularis) | 214 | 72 | 33.645 |
| Lexicality by Morphology | 3 | Left angular gyrus | 188 | 142 | 75.532 |
| Lexicality by Morphology | 4 | Right cerebellum crus I | 57 | 0 | 0 |
| Lexicality by Morphology | 5 | Left middle temporal gyrus | 53 | 41 | 77.358 |

**Supplementary Table S3.**
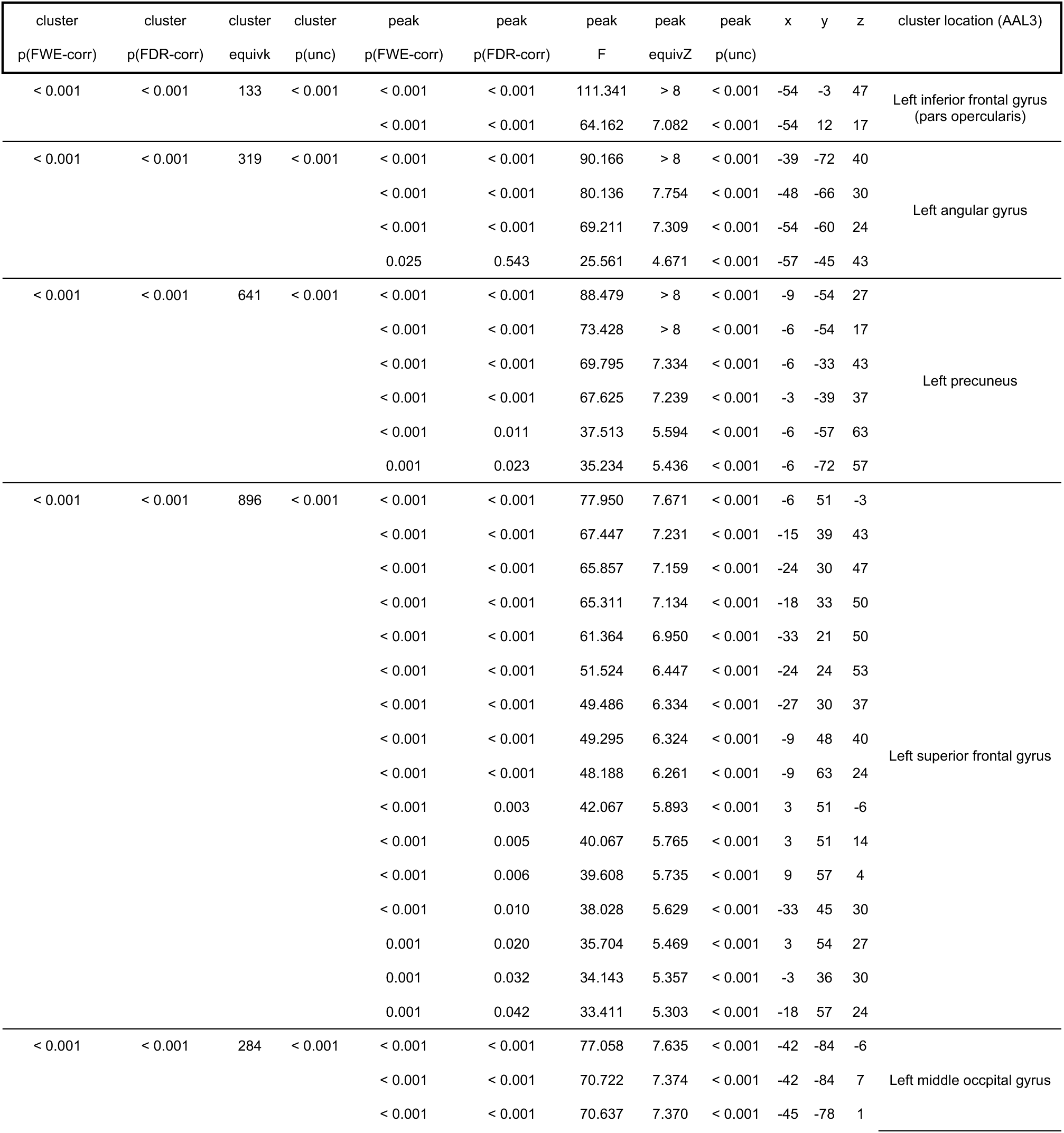
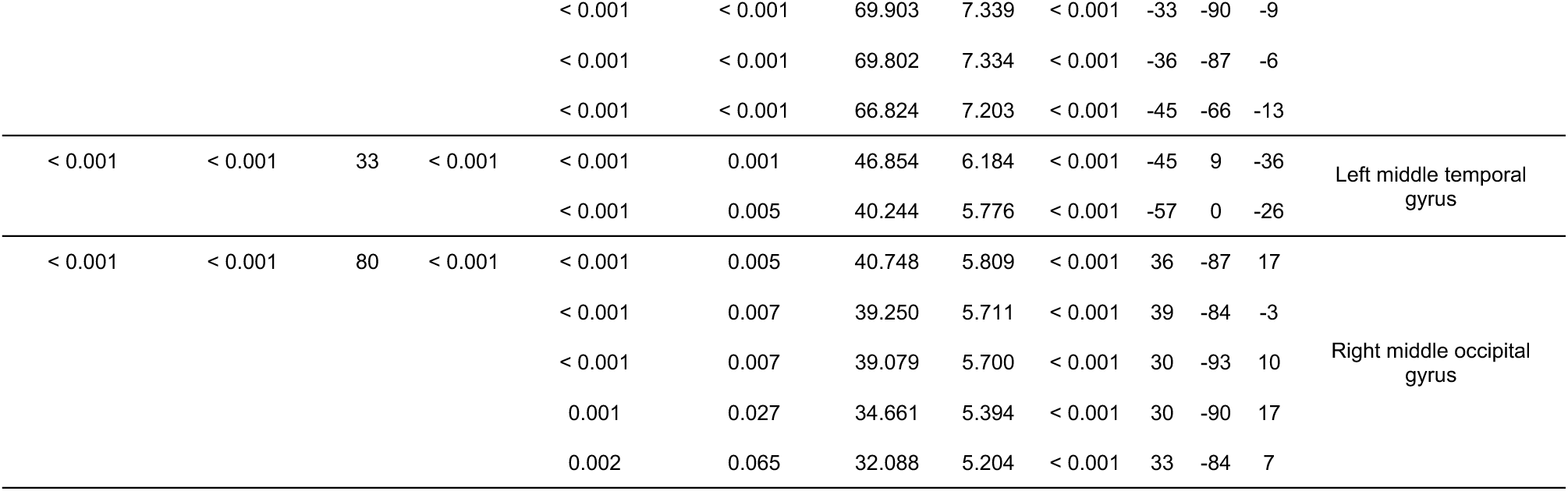
| Main effect of Lexicality in univariate fMRI data. Coordinates in the standard Montreal Neurological Institute (MNI) space of significant (p< 0.05 FWE-corrected, k>30) effects and cluster localization according to the AAL3 template^3^. The term “k” indicates the total number of voxels in each cluster.

| cluster<br>p(FWE-corr) | cluster<br>p(FDR-corr) | cluster<br>equivk | cluster<br>p(unc) | peak<br>p(FWE-corr) | peak<br>p(FDR-corr) | peak<br>F | peak<br>equivZ | peak<br>p(unc) | x | y | z | cluster location (AAL3) |
| --- | --- | --- | --- | --- | --- | --- | --- | --- | --- | --- | --- | --- |
| < 0.001 | < 0.001 | 133 | < 0.001 | < 0.001 | < 0.001 | 111.341 | > 8 | < 0.001 | -54 | -3 | 47 | Left inferior frontal gyrus<br>(pars opercularis) |
|  |  |  |  | < 0.001 | < 0.001 | 64.162 | 7.082 | < 0.001 | -54 | 12 | 17 |  |
| < 0.001 | < 0.001 | 319 | < 0.001 | < 0.001 | < 0.001 | 90.166 | > 8 | < 0.001 | -39 | -72 | 40 | Left angular gyrus |
|  |  |  |  | < 0.001 | < 0.001 | 80.136 | 7.754 | < 0.001 | -48 | -66 | 30 |  |
|  |  |  |  | < 0.001 | < 0.001 | 69.211 | 7.309 | < 0.001 | -54 | -60 | 24 |  |
|  |  |  |  | 0.025 | 0.543 | 25.561 | 4.671 | < 0.001 | -57 | -45 | 43 |  |
| < 0.001 | < 0.001 | 641 | < 0.001 | < 0.001 | < 0.001 | 88.479 | > 8 | < 0.001 | -9 | -54 | 27 | Left precuneus |
|  |  |  |  | < 0.001 | < 0.001 | 73.428 | > 8 | < 0.001 | -6 | -54 | 17 |  |
|  |  |  |  | < 0.001 | < 0.001 | 69.795 | 7.334 | < 0.001 | -6 | -33 | 43 |  |
|  |  |  |  | < 0.001 | < 0.001 | 67.625 | 7.239 | < 0.001 | -3 | -39 | 37 |  |
|  |  |  |  | < 0.001 | 0.011 | 37.513 | 5.594 | < 0.001 | -6 | -57 | 63 |  |
|  |  |  |  | 0.001 | 0.023 | 35.234 | 5.436 | < 0.001 | -6 | -72 | 57 |  |
| < 0.001 | < 0.001 | 896 | < 0.001 | < 0.001 | < 0.001 | 77.950 | 7.671 | < 0.001 | -6 | 51 | -3 | Left superior frontal gyrus |
|  |  |  |  | < 0.001 | < 0.001 | 67.447 | 7.231 | < 0.001 | -15 | 39 | 43 |  |
|  |  |  |  | < 0.001 | < 0.001 | 65.857 | 7.159 | < 0.001 | -24 | 30 | 47 |  |
|  |  |  |  | < 0.001 | < 0.001 | 65.311 | 7.134 | < 0.001 | -18 | 33 | 50 |  |
|  |  |  |  | < 0.001 | < 0.001 | 61.364 | 6.950 | < 0.001 | -33 | 21 | 50 |  |
|  |  |  |  | < 0.001 | < 0.001 | 51.524 | 6.447 | < 0.001 | -24 | 24 | 53 |  |
|  |  |  |  | < 0.001 | < 0.001 | 49.486 | 6.334 | < 0.001 | -27 | 30 | 37 |  |
|  |  |  |  | < 0.001 | < 0.001 | 49.295 | 6.324 | < 0.001 | -9 | 48 | 40 |  |
|  |  |  |  | < 0.001 | < 0.001 | 48.188 | 6.261 | < 0.001 | -9 | 63 | 24 |  |
|  |  |  |  | < 0.001 | 0.003 | 42.067 | 5.893 | < 0.001 | 3 | 51 | -6 |  |
|  |  |  |  | < 0.001 | 0.005 | 40.067 | 5.765 | < 0.001 | 3 | 51 | 14 |  |
|  |  |  |  | < 0.001 | 0.006 | 39.608 | 5.735 | < 0.001 | 9 | 57 | 4 |  |
|  |  |  |  | < 0.001 | 0.010 | 38.028 | 5.629 | < 0.001 | -33 | 45 | 30 |  |
|  |  |  |  | 0.001 | 0.020 | 35.704 | 5.469 | < 0.001 | 3 | 54 | 27 |  |
|  |  |  |  | 0.001 | 0.032 | 34.143 | 5.357 | < 0.001 | -3 | 36 | 30 |  |
|  |  |  |  | 0.001 | 0.042 | 33.411 | 5.303 | < 0.001 | -18 | 57 | 24 |  |
| < 0.001 | < 0.001 | 284 | < 0.001 | < 0.001 | < 0.001 | 77.058 | 7.635 | < 0.001 | -42 | -84 | -6 | Left middle occipital gyrus |
|  |  |  |  | < 0.001 | < 0.001 | 70.722 | 7.374 | < 0.001 | -42 | -84 | 7 |  |
|  |  |  |  | < 0.001 | < 0.001 | 70.637 | 7.370 | < 0.001 | -45 | -78 | 1 |  |
|  |  |  |  | < 0.001 | < 0.001 | 69.903 | 7.339 | < 0.001 | -33 | -90 | -9 |  |
|  |  |  |  | < 0.001 | < 0.001 | 69.802 | 7.334 | < 0.001 | -36 | -87 | -6 |  |
|  |  |  |  | < 0.001 | < 0.001 | 66.824 | 7.203 | < 0.001 | -45 | -66 | -13 |  |
| < 0.001 | < 0.001 | 33 | < 0.001 | < 0.001 | 0.001 | 46.854 | 6.184 | < 0.001 | -45 | 9 | -36 | Left middle temporal gyrus |
|  |  |  |  | < 0.001 | 0.005 | 40.244 | 5.776 | < 0.001 | -57 | 0 | -26 |  |
| < 0.001 | < 0.001 | 80 | < 0.001 | < 0.001 | 0.005 | 40.748 | 5.809 | < 0.001 | 36 | -87 | 17 | Right middle occipital gyrus |
|  |  |  |  | < 0.001 | 0.007 | 39.250 | 5.711 | < 0.001 | 39 | -84 | -3 |  |
|  |  |  |  | < 0.001 | 0.007 | 39.079 | 5.700 | < 0.001 | 30 | -93 | 10 |  |
|  |  |  |  | 0.001 | 0.027 | 34.661 | 5.394 | < 0.001 | 30 | -90 | 17 |  |
|  |  |  |  | 0.002 | 0.065 | 32.088 | 5.204 | < 0.001 | 33 | -84 | 7 |  |

**Supplementary Table S4.** | Results of the Existing > Novel Words contrast in univariate fMRI data. Coordinates in the standard MNI space of significant (p< 0.05 FWE-corrected, k>30) effects and cluster localization according to the AAL3 template^3^. The term “k” indicates the total number of voxels in each cluster.

| cluster<br>p(FWE-corr) | cluster<br>p(FDR-corr) | cluster<br>equivk | cluster<br>p(unc) | peak<br>p(FWE-corr) | peak<br>p(FDR-corr) | peak<br>T | peak<br>equivZ | peak<br>p(unc) | x | y | z | cluster location (AAL3) |
| --- | --- | --- | --- | --- | --- | --- | --- | --- | --- | --- | --- | --- |
| < 0.001 | 0.000 | 343 | < 0.001 | < 0.001 | < 0.001 | 9.496 | >8 | < 0.001 | -39 | -72 | 40 | Left angular gyrus |
|  |  |  |  | < 0.001 | < 0.001 | 8.952 | 7.836 | < 0.001 | -48 | -66 | 30 |  |
|  |  |  |  | < 0.001 | < 0.001 | 8.319 | 7.401 | < 0.001 | -54 | -60 | 24 |  |
|  |  |  |  | 0.013 | 0.333 | 5.056 | 4.811 | < 0.001 | -57 | -45 | 43 |  |
| < 0.001 | 0.000 | 706 | < 0.001 | < 0.001 | < 0.001 | 9.406 | >8 | < 0.001 | -9 | -54 | 27 | Left precuneus |
|  |  |  |  | < 0.001 | < 0.001 | 8.569 | 7.578 | < 0.001 | -6 | -54 | 17 |  |
|  |  |  |  | < 0.001 | < 0.001 | 8.354 | 7.426 | < 0.001 | -6 | -33 | 43 |  |
|  |  |  |  | < 0.001 | < 0.001 | 8.223 | 7.332 | < 0.001 | -3 | -39 | 37 |  |
|  |  |  |  | < 0.001 | 0.007 | 6.125 | 5.714 | < 0.001 | -6 | -57 | 63 |  |
|  |  |  |  | < 0.001 | 0.013 | 5.936 | 5.558 | < 0.001 | -6 | -72 | 57 |  |
|  |  |  |  | 0.014 | 0.352 | 5.018 | 4.779 | < 0.001 | -12 | -54 | 50 |  |
| < 0.001 | 0.000 | 989 | < 0.001 | < 0.001 | < 0.001 | 8.829 | 7.754 | < 0.001 | -6 | 51 | -3 | Left superior frontal gyrus |
|  |  |  |  | < 0.001 | < 0.001 | 8.213 | 7.324 | < 0.001 | -15 | 39 | 43 |  |
|  |  |  |  | < 0.001 | < 0.001 | 8.115 | 7.254 | < 0.001 | -24 | 30 | 47 |  |
|  |  |  |  | < 0.001 | < 0.001 | 8.082 | 7.229 | < 0.001 | -18 | 33 | 50 |  |
|  |  |  |  | < 0.001 | < 0.001 | 7.834 | 7.047 | < 0.001 | -33 | 21 | 50 |  |
|  |  |  |  | < 0.001 | < 0.001 | 7.178 | 6.551 | < 0.001 | -24 | 24 | 53 |  |
|  |  |  |  | < 0.001 | < 0.001 | 7.035 | 6.440 | < 0.001 | -27 | 30 | 37 |  |
|  |  |  |  | < 0.001 | < 0.001 | 7.021 | 6.430 | < 0.001 | -9 | 48 | 40 |  |
|  |  |  |  | < 0.001 | < 0.001 | 6.942 | 6.368 | < 0.001 | -9 | 63 | 24 |  |
|  |  |  |  | < 0.001 | 0.002 | 6.486 | 6.007 | < 0.001 | 3 | 51 | -6 |  |
|  |  |  |  | < 0.001 | 0.003 | 6.330 | 5.881 | < 0.001 | 3 | 51 | 14 |  |
|  |  |  |  | < 0.001 | 0.004 | 6.293 | 5.851 | < 0.001 | 9 | 57 | 4 |  |
|  |  |  |  | < 0.001 | 0.006 | 6.167 | 5.748 | < 0.001 | -33 | 45 | 30 |  |
|  |  |  |  | < 0.001 | 0.012 | 5.975 | 5.590 | < 0.001 | 3 | 54 | 27 |  |
|  |  |  |  | < 0.001 | 0.019 | 5.843 | 5.481 | < 0.001 | -3 | 36 | 30 |  |
|  |  |  |  | 0.001 | 0.024 | 5.780 | 5.428 | < 0.001 | -18 | 57 | 24 |  |
| < 0.001 | 0.002 | 38 | < 0.001 | < 0.001 | < 0.001 | 6.845 | 6.292 | < 0.001 | -45 | 9 | -36 | Left middle temporal gyrus |
|  |  |  |  | < 0.001 | 0.003 | 6.344 | 5.892 | < 0.001 | -57 | 0 | -26 |  |
| < 0.001 | 0.003 | 33 | 0.001 | < 0.001 | 0.007 | 6.093 | 5.687 | < 0.001 | 54 | -36 | 34 | Right supramarginal gyrus |
|  |  |  |  | < 0.001 | 0.008 | 6.075 | 5.672 | < 0.001 | 57 | -36 | 27 |  |

**Supplementary Table S5.** | Results of the Novel > Existing Words contrast in univariate fMRI data. Coordinates in the standard MNI space of significant (p< 0.05 FWE-corrected, k>30) effects and cluster localization according to the AAL3 template^3^. The term “k” indicates the total number of voxels in each cluster.

| cluster<br>p(FWE-corr) | cluster<br>p(FDR-corr) | cluster<br>equivk | cluster<br>p(unc) | peak<br>p(FWE-corr) | peak<br>p(FDR-corr) | peak<br>T | peak<br>equivZ | peak<br>p(unc) | x | y | z | cluster location (AAL3) |
| --- | --- | --- | --- | --- | --- | --- | --- | --- | --- | --- | --- | --- |
| < 0.001 | 0.000 | 145 | < 0.001 | < 0.001 | < 0.001 | 10.552 | > 8 | < 0.001 | -54 | -3 | 47 | Left inferior frontal gyrus<br>(pars opercularis) |
|  |  |  |  | < 0.001 | < 0.001 | 8.010 | 7.177 | < 0.001 | -54 | 12 | 17 |  |
| < 0.001 | 0.000 | 301 | < 0.001 | < 0.001 | < 0.001 | 8.778 | 7.721 | < 0.001 | -42 | -84 | -6 | Left middle occipital gyrus |
|  |  |  |  | < 0.001 | < 0.001 | 8.410 | 7.465 | < 0.001 | -42 | -84 | 7 |  |
|  |  |  |  | < 0.001 | < 0.001 | 8.405 | 7.462 | < 0.001 | -45 | -78 | 1 |  |
|  |  |  |  | < 0.001 | < 0.001 | 8.361 | 7.431 | < 0.001 | -33 | -90 | -9 |  |
|  |  |  |  | < 0.001 | < 0.001 | 8.355 | 7.426 | < 0.001 | -36 | -87 | -6 |  |
|  |  |  |  | < 0.001 | < 0.001 | 8.175 | 7.297 | < 0.001 | -45 | -66 | -13 |  |
| < 0.001 | 0.000 | 98 | < 0.001 | < 0.001 | 0.002 | 6.383 | 5.924 | < 0.001 | 36 | -87 | 17 | Right middle occipital gyrus |
|  |  |  |  | < 0.001 | 0.003 | 6.265 | 5.828 | < 0.001 | 39 | -84 | -3 |  |
|  |  |  |  | < 0.001 | 0.003 | 6.251 | 5.817 | < 0.001 | 30 | -93 | 10 |  |
|  |  |  |  | < 0.001 | 0.014 | 5.887 | 5.518 | < 0.001 | 30 | -90 | 17 |  |
|  |  |  |  | 0.001 | 0.033 | 5.665 | 5.331 | < 0.001 | 33 | -84 | 7 |  |

**Supplementary Table S6.**
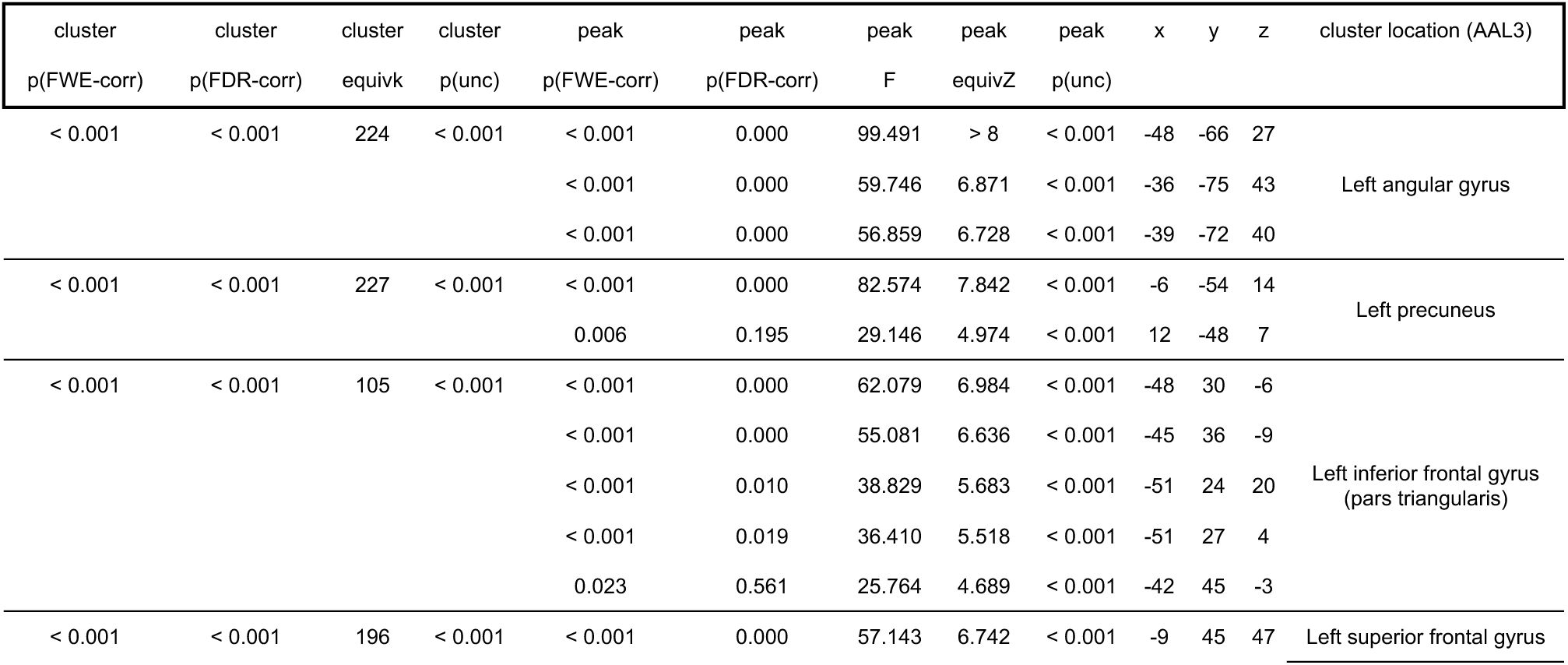
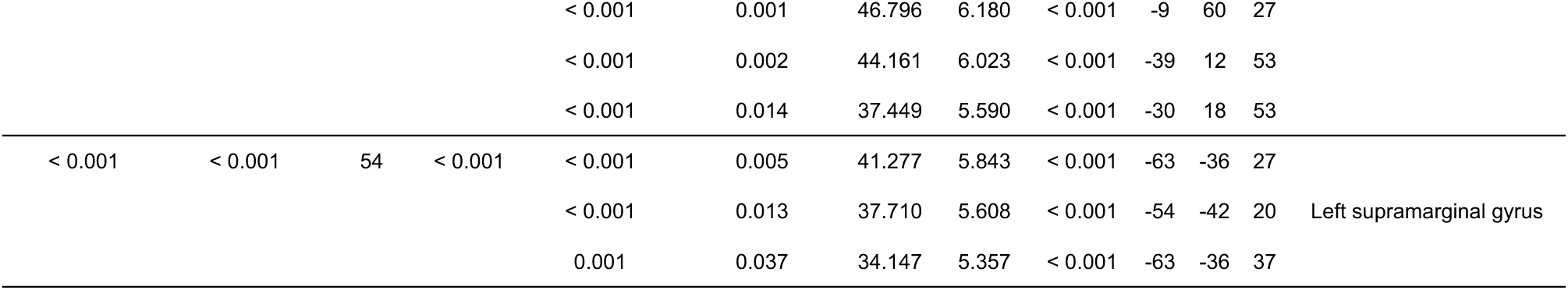
| Main effect of Morphology in univariate fMRI data. Coordinates in the standard MNI space of significant (p< 0.05 FWE-corrected, k>30) effects and cluster localization according to the AAL3 template^3^. The term “k” indicates the total number of voxels in each cluster.

**Supplementary Table S7.**
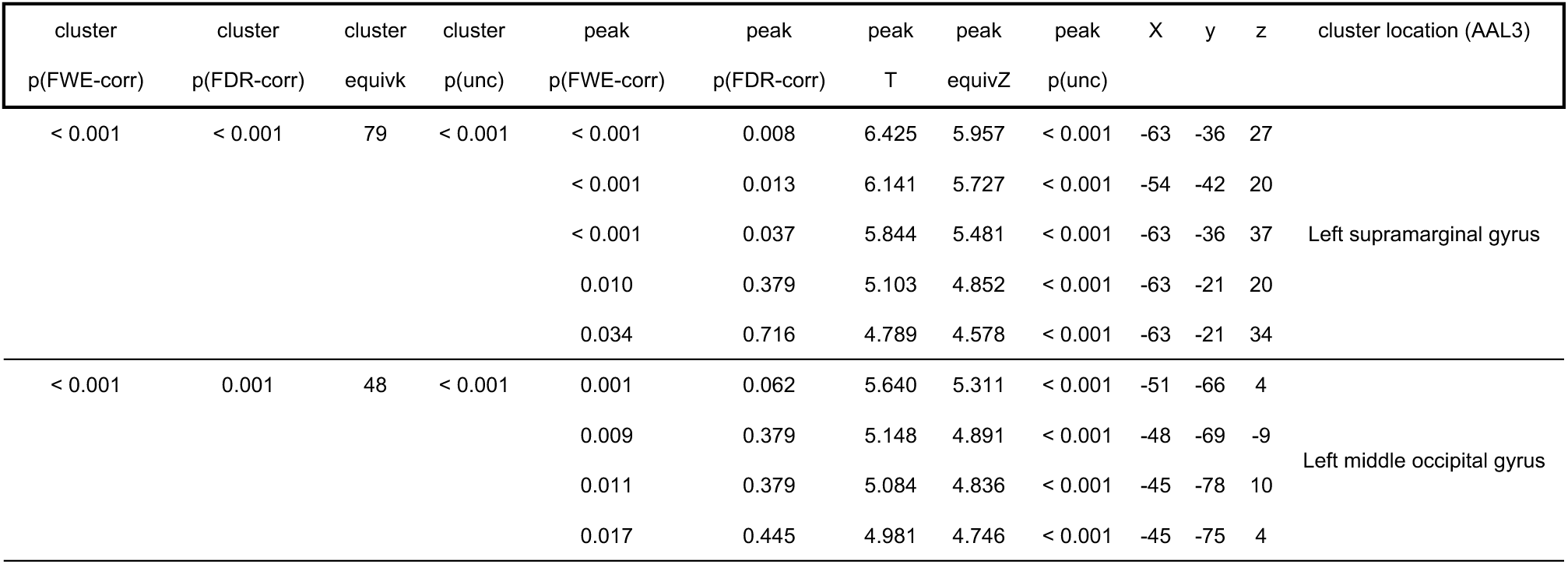
| Results of the Simple > Complex contrast in univariate fMRI data. Coordinates in the standard MNI space of significant (p< 0.05 FWE-corrected, k>30) effects and cluster localization according to the AAL3 template^3^. The term “k” indicates the total number of voxels in each cluster.

**Supplementary Table S8.**
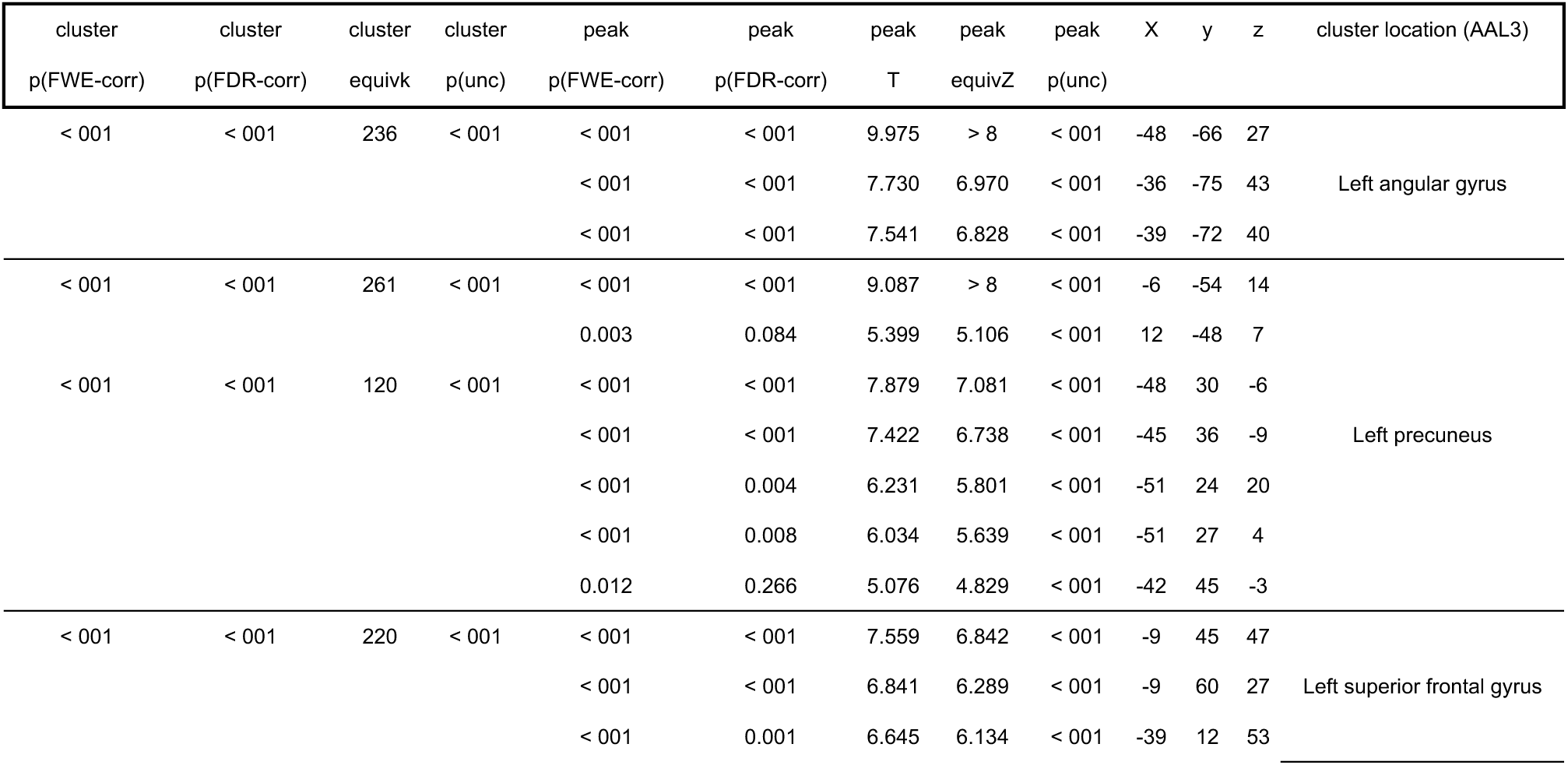
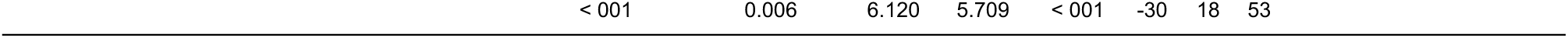
| Results of the Complex > Simple contrast in univariate fMRI data. Coordinates in the standard MNI space of significant (p< 0.05 FWE-corrected, k>30) effects and cluster localization according to the AAL3 template^3^. The term “k” indicates the total number of voxels in each cluster.

**Supplementary Table S9.** | Lexicality-by-Morphology interaction in univariate fMRI data. Coordinates in the standard Montreal Neurological Institute (MNI) space of significant (p< 0.05 FWE-corrected, k>30) effects and cluster localization according to the AAL3 template^3^. The term “k” indicates the total number of voxels in each cluster.

| cluster<br>p(FWE-corr) | cluster<br>p(FDR-corr) | cluster<br>equivk | cluster<br>p(unc) | peak<br>p(FWE-corr) | peak<br>p(FDR-corr) | peak<br>F | peak<br>equivZ | peak<br>p(unc) | x | y | z | cluster location (AAL3) |
| --- | --- | --- | --- | --- | --- | --- | --- | --- | --- | --- | --- | --- |
| < 0.001 | < 0.001 | 214 | < 0.001 | < 0.001 | < 0.001 | 89.411 | > 8 | < 0.001 | -48 | 30 | -9 | Left inferior frontal gyrus<br>(pars triangularis) |
|  |  |  |  | < 0.001 | 0.001 | 46.968 | 6.190 | < 0.001 | -54 | 27 | 4 |  |
|  |  |  |  | < 0.001 | 0.001 | 46.083 | 6.138 | < 0.001 | -54 | 24 | 17 |  |
|  |  |  |  | 0.001 | 0.038 | 33.045 | 5.276 | < 0.001 | -51 | 36 | 7 |  |
| < 0.001 | < 0.001 | 396 | < 0.001 | < 0.001 | < 0.001 | 85.444 | > 8 | < 0.001 | -12 | 33 | 53 | Left medial superior frontal<br>gyrus |
|  |  |  |  | < 0.001 | < 0.001 | 66.632 | 7.194 | < 0.001 | -3 | 42 | 47 |  |
|  |  |  |  | < 0.001 | < 0.001 | 64.546 | 7.099 | < 0.001 | -36 | 12 | 53 |  |
|  |  |  |  | < 0.001 | < 0.001 | 64.237 | 7.085 | < 0.001 | -33 | 15 | 57 |  |
|  |  |  |  | < 0.001 | < 0.001 | 58.248 | 6.798 | < 0.001 | -12 | 45 | 47 |  |
|  |  |  |  | < 0.001 | < 0.001 | 56.227 | 6.695 | < 0.001 | -12 | 57 | 30 |  |
|  |  |  |  | < 0.001 | < 0.001 | 53.795 | 6.569 | < 0.001 | -12 | 51 | 37 |  |
|  |  |  |  | < 0.001 | 0.007 | 38.239 | 5.644 | < 0.001 | -24 | 24 | 53 |  |
| < 0.001 | < 0.001 | 53 | < 0.001 | < 0.001 | < 0.001 | 78.759 | 7.701 | < 0.001 | -63 | -45 | -3 | Left middle temporal gyrus |
| < 0.001 | < 0.001 | 57 | < 0.001 | < 0.001 | 0.001 | 47.006 | 6.192 | < 0.001 | 9 | -81 | -29 | Right cerebellum crus I |
|  |  |  |  | < 0.001 | 0.001 | 45.635 | 6.112 | < 0.001 | 30 | -69 | -32 |  |
|  |  |  |  | 0.001 | 0.038 | 33.186 | 5.287 | < 0.001 | 30 | -75 | -29 |  |
| < 0.001 | < 0.001 | 188 | < 0.001 | < 0.001 | 0.002 | 42.709 | 5.933 | < 0.001 | -42 | -60 | 47 | Left angular gyrus |
|  |  |  |  | < 0.001 | 0.003 | 41.084 | 5.831 | < 0.001 | -45 | -57 | 37 |  |
|  |  |  |  | < 0.001 | 0.003 | 40.840 | 5.815 | < 0.001 | -45 | -54 | 50 |  |
|  |  |  |  | < 0.001 | 0.009 | 37.435 | 5.589 | < 0.001 | -45 | -48 | 47 |  |
|  |  |  |  | 0.001 | 0.033 | 33.815 | 5.333 | < 0.001 | -54 | -57 | 24 |  |
|  |  |  |  | 0.002 | 0.055 | 31.909 | 5.191 | < 0.001 | -33 | -72 | 50 |  |
|  |  |  |  | 0.003 | 0.065 | 31.375 | 5.150 | < 0.001 | -45 | -66 | 30 |  |

**Supplementary Table S10.** | Results of the Existing Words > Novel Words contrast in RSA. Coordinates in the standard MNI space of significant (p< 0.001 uncorrected, k>0) effects and cluster localization according to the AAL3 template^3^. Coordinates highlighted in bold are significant at a p< 0.05 FWE-corrected threshold. The term “k” indicates the total number of voxels in each cluster.

| cluster<br>p(FWE-corr) | cluster<br>p(FDR-corr) | cluster<br>equivk | cluster<br>p(unc) | peak<br>p(FWE-corr) | peak<br>p(FDR-corr) | peak<br>T | peak<br>equivZ | peak<br>p(unc) | x | y | z | cluster location (AAL3) |
| --- | --- | --- | --- | --- | --- | --- | --- | --- | --- | --- | --- | --- |
| < 0.001 | 0.002 | 46 | < 0.001 | **< 0.001** | **0.011** | **5.717** | **5.375** | **< 0.001** | **-6** | **-54** | **27** | Left precuneus |
|  |  |  |  | **0.049** | **0.106** | **4.275** | **4.121** | **< 0.001** | **-12** | **-54** | **30** |  |
|  |  |  |  | 0.053 | 0.106 | 4.257 | 4.104 | < 0.001 | -12 | -54 | 17 |  |
| 0.001 | 0.006 | 33 | < 0.001 | **0.009** | **0.038** | **4.702** | **4.502** | **< 0.001** | **-54** | **0** | **-29** | Left middle temporal gyrus |
|  |  |  |  | **0.011** | **0.038** | **4.653** | **4.459** | **< 0.001** | **-51** | **3** | **-36** |  |
|  |  |  |  | **0.011** | **0.038** | **4.653** | **4.459** | **< 0.001** | **-48** | **6** | **-36** |  |
|  |  |  |  | **0.011** | **0.038** | **4.653** | **4.459** | **< 0.001** | **-45** | **9** | **-36** |  |
|  |  |  |  | **0.011** | **0.038** | **4.653** | **4.459** | **< 0.001** | **-48** | **0** | **-32** |  |
|  |  |  |  | **0.011** | **0.038** | **4.653** | **4.459** | **< 0.001** | **-51** | **6** | **-32** |  |
|  |  |  |  | **0.011** | **0.038** | **4.653** | **4.459** | **< 0.001** | **-45** | **6** | **-32** |  |
|  |  |  |  | **0.011** | **0.038** | **4.653** | **4.459** | **< 0.001** | **-48** | **9** | **-32** |  |
|  |  |  |  | **0.011** | **0.038** | **4.653** | **4.459** | **< 0.001** | **-48** | **3** | **-29** |  |
|  |  |  |  | **0.019** | **0.051** | **4.509** | **4.330** | **< 0.001** | **-57** | **-3** | **-23** |  |
|  |  |  |  | **0.019** | **0.051** | **4.509** | **4.330** | **< 0.001** | **-54** | **0** | **-23** |  |
|  |  |  |  | **0.019** | **0.051** | **4.509** | **4.330** | **< 0.001** | **-57** | **0** | **-19** |  |
| < 0.001 | 0.004 | 37 | < 0.001 | **0.036** | **0.083** | **4.358** | **4.196** | **< 0.001** | **-54** | **-54** | **40** | Left inferior parietal gyrus |
|  |  |  |  | 0.052 | 0.106 | 4.263 | 4.110 | < 0.001 | -48 | -54 | 47 |  |
|  |  |  |  | 0.062 | 0.117 | 4.216 | 4.068 | < 0.001 | -57 | -45 | 43 |  |
|  |  |  |  | 0.192 | 0.254 | 3.886 | 3.768 | < 0.001 | -54 | -60 | 40 |  |
| 0.105 | 0.260 | 7 | 0.065 | 0.070 | 0.125 | 4.186 | 4.040 | < 0.001 | 0 | -24 | 37 | Left middle cingulum |
|  |  |  |  | 0.714 | 0.844 | 3.275 | 3.201 | < 0.001 | -6 | -21 | 43 |  |
| 0.002 | 0.011 | 27 | 0.001 | 0.093 | 0.156 | 4.109 | 3.971 | < 0.001 | -21 | 21 | 47 | Left superior frontal gyrus |
|  |  |  |  | 0.127 | 0.192 | 4.027 | 3.896 | < 0.001 | -24 | 33 | 34 |  |
|  |  |  |  | 0.132 | 0.193 | 4.016 | 3.886 | < 0.001 | -24 | 21 | 40 |  |
|  |  |  |  | 0.218 | 0.279 | 3.841 | 3.726 | < 0.001 | -27 | 24 | 47 |  |
|  |  |  |  | 0.280 | 0.347 | 3.745 | 3.638 | < 0.001 | -24 | 39 | 34 |  |
|  |  |  |  | 0.547 | 0.604 | 3.443 | 3.358 | < 0.001 | -33 | 30 | 37 |  |
| 0.228 | 0.304 | 4 | 0.152 | 0.119 | 0.187 | 4.045 | 3.913 | < 0.001 | -57 | -57 | 30 | Left supramarginal gyrus |
| 0.174 | 0.270 | 5 | 0.113 | 0.147 | 0.212 | 3.978 | 3.851 | < 0.001 | -42 | -63 | -3 | Left middle temporal gyrus |
|  |  |  |  | 0.509 | 0.571 | 3.481 | 3.394 | < 0.001 | -48 | -60 | -6 |  |
| 0.022 | 0.094 | 14 | 0.013 | 0.157 | 0.220 | 3.957 | 3.832 | < 0.001 | -18 | 42 | 47 | Left superior frontal gyrus |
|  |  |  |  | 0.426 | 0.478 | 3.568 | 3.474 | < 0.001 | -21 | 45 | 40 |  |
| 0.033 | 0.120 | 12 | 0.020 | 0.166 | 0.228 | 3.937 | 3.814 | < 0.001 | -42 | -75 | 34 | Left middle occipital gyrus |
|  |  |  |  | 0.242 | 0.307 | 3.801 | 3.689 | < 0.001 | -36 | -72 | 37 |  |
|  |  |  |  | 0.508 | 0.571 | 3.482 | 3.394 | < 0.001 | -39 | -75 | 40 |  |
| 0.082 | 0.227 | 8 | 0.050 | 0.186 | 0.251 | 3.898 | 3.778 | < 0.001 | -21 | 18 | 57 | Left superior frontal gyrus |
|  |  |  |  | 0.375 | 0.421 | 3.625 | 3.528 | < 0.001 | -24 | 18 | 63 |  |
| 0.228 | 0.304 | 4 | 0.152 | 0.216 | 0.279 | 3.844 | 3.729 | < 0.001 | 3 | 51 | 30 | Right medial superior frontal gyrus |
|  |  |  |  | 0.319 | 0.389 | 3.693 | 3.590 | < 0.001 | 6 | 48 | 27 |  |
| 0.174 | 0.270 | 5 | 0.113 | 0.247 | 0.307 | 3.794 | 3.683 | < 0.001 | -42 | -57 | 24 | Left angular gyrus |
| 0.065 | 0.203 | 9 | 0.039 | 0.316 | 0.389 | 3.698 | 3.594 | < 0.001 | -6 | -54 | 63 | Left precuneus |
| 0.174 | 0.270 | 5 | 0.113 | 0.343 | 0.409 | 3.663 | 3.562 | < 0.001 | -48 | -51 | -16 | Left inferior temporal gyrus |
|  |  |  |  | 0.767 | 0.915 | 3.217 | 3.147 | < 0.001 | -45 | -48 | -13 |  |
| 0.228 | 0.304 | 4 | 0.152 | 0.344 | 0.409 | 3.662 | 3.561 | < 0.001 | -6 | -39 | 40 | Left middle cingulum |
| 0.302 | 0.401 | 3 | 0.211 | 0.372 | 0.421 | 3.629 | 3.531 | < 0.001 | -42 | -84 | 4 | Left middle occipital gyrus |
| 0.174 | 0.270 | 5 | 0.113 | 0.375 | 0.421 | 3.626 | 3.528 | < 0.001 | -3 | -60 | 40 | Left precuneus |
| 0.134 | 0.270 | 6 | 0.085 | 0.375 | 0.421 | 3.625 | 3.527 | < 0.001 | -36 | 18 | 47 | Left middle frontal gyrus |
|  |  |  |  | 0.706 | 0.838 | 3.283 | 3.208 | < 0.001 | -36 | 18 | 57 |  |
| 0.553 | 0.474 | 1 | 0.474 | 0.415 | 0.470 | 3.581 | 3.486 | < 0.001 | -18 | 39 | 34 | Left superior frontal gyrus |
| 0.174 | 0.270 | 5 | 0.113 | 0.459 | 0.520 | 3.533 | 3.442 | < 0.001 | -6 | 42 | 47 | Left medial superior frontal gyrus |
|  |  |  |  | 0.466 | 0.521 | 3.526 | 3.435 | < 0.001 | -9 | 45 | 50 |  |
| 0.553 | 0.474 | 1 | 0.474 | 0.538 | 0.604 | 3.453 | 3.367 | < 0.001 | -6 | 57 | 7 | Left medial superior frontal gyrus |
| 0.553 | 0.474 | 1 | 0.474 | 0.541 | 0.604 | 3.449 | 3.363 | < 0.001 | -39 | 12 | 53 | Left middle frontal gyrus |
| 0.553 | 0.474 | 1 | 0.474 | 0.578 | 0.647 | 3.412 | 3.330 | < 0.001 | -39 | 21 | 43 | Left middle frontal gyrus |
| 0.553 | 0.474 | 1 | 0.474 | 0.659 | 0.780 | 3.332 | 3.254 | < 0.001 | -9 | -60 | 63 | Left precuneus |
| 0.553 | 0.474 | 1 | 0.474 | 0.665 | 0.780 | 3.325 | 3.248 | < 0.001 | -24 | 15 | 50 | Left superior frontal gyrus |
| 0.405 | 0.474 | 2 | 0.306 | 0.668 | 0.780 | 3.323 | 3.246 | < 0.001 | -9 | 36 | 50 | Left medial superior frontal gyrus |
| 0.553 | 0.474 | 1 | 0.474 | 0.669 | 0.780 | 3.321 | 3.244 | < 0.001 | -9 | 60 | 4 | Left medial superior frontal gyrus |
| 0.405 | 0.474 | 2 | 0.306 | 0.680 | 0.792 | 3.310 | 3.233 | < 0.001 | -15 | -27 | 40 | NA: outside template |
| 0.553 | 0.474 | 1 | 0.474 | 0.735 | 0.883 | 3.252 | 3.180 | < 0.001 | -3 | 45 | 20 | Left medial superior frontal gyrus |
| 0.553 | 0.474 | 1 | 0.474 | 0.759 | 0.915 | 3.226 | 3.155 | < 0.001 | 3 | 54 | 7 | NA: outside template |
| 0.405 | 0.474 | 2 | 0.306 | 0.759 | 0.915 | 3.226 | 3.155 | < 0.001 | -51 | -66 | 37 | Left angular gyrus |
| 0.553 | 0.474 | 1 | 0.474 | 0.774 | 0.921 | 3.209 | 3.139 | < 0.001 | -24 | 27 | 57 | Left middle frontal gyrus |
| 0.553 | 0.474 | 1 | 0.474 | 0.780 | 0.927 | 3.202 | 3.132 | < 0.001 | -30 | 42 | 27 | Left middle frontal gyrus |
| 0.553 | 0.474 | 1 | 0.474 | 0.789 | 0.938 | 3.192 | 3.123 | < 0.001 | -27 | 15 | 57 | Left middle frontal gyrus |
| 0.553 | 0.474 | 1 | 0.474 | 0.808 | 0.978 | 3.169 | 3.101 | < 0.001 | -33 | -78 | 37 | Left middle occipital gyrus |
| 0.553 | 0.474 | 1 | 0.474 | 0.810 | 0.978 | 3.166 | 3.099 | < 0.001 | -3 | -57 | 60 | Left precuneus |

**Supplementary Table S11.** | RSA results for Morphologically Simple Existing Words. Coordinate in the standard MNI space of the significant (p< 0.001 uncorrected, k>0) effect, and cluster localization according to the AAL3 template^3^. The term “k” indicates the total number of voxels in each cluster.

| cluster | cluster | cluster | cluster | peak | peak | peak | peak | peak | x | y | z | cluster location (AAL3) |
| --- | --- | --- | --- | --- | --- | --- | --- | --- | --- | --- | --- | --- |
| p(FWE-corr) | p(FDR-corr) | equivk | p(unc) | p(FWE-corr) | p(FDR-corr) | T | equivZ | p(unc) |  |  |  |  |
| 0.553 | 0.474 | 1 | 0.474 | 0.698 | 0.705 | 3.291 | 3.216 | < 0.001 | -6 | -54 | 27 | Left precuneus |

**Supplementary Table S12.** | Results of the Morphologically Complex > Simple contrast in RSA. Coordinates in the standard MNI space of significant (p< 0.001 uncorrected, k>0) effects and cluster localization according to the AAL3 template^3^. Coordinates highlighted in bold are significant at a p< 0.05 FWE-corrected threshold. The term “k” indicates the total number of voxels in each cluster.

| cluster | cluster | cluster | cluster | peak | peak | peak | peak | peak | x | y | z | cluster location (AAL3) |
| --- | --- | --- | --- | --- | --- | --- | --- | --- | --- | --- | --- | --- |
| p(FWE-corr) | p(FDR-corr) | equivk | p(unc) | p(FWE-corr) | p(FDR-corr) | T | equivZ | p(unc) |  |  |  |  |
| < 0.001 | < 0.001 | 93 | < 0.001 | **< 0.001** | **0.003** | **5.789** | **5.435** | **< 0.001** | **-18** | **33** | **40** | Left superior frontal gyrus |
|  |  |  |  | **0.001** | **0.016** | **5.209** | **4.944** | **< 0.001** | **-18** | **39** | **47** |  |
|  |  |  |  | **0.007** | **0.061** | **4.758** | **4.551** | **< 0.001** | **-21** | **42** | **43** |  |
|  |  |  |  | 0.065 | 0.127 | 4.204 | 4.057 | < 0.001 | -21 | 21 | 53 |  |
|  |  |  |  | 0.079 | 0.127 | 4.155 | 4.012 | < 0.001 | -15 | 42 | 40 |  |
|  |  |  |  | 0.172 | 0.190 | 3.925 | 3.803 | < 0.001 | -30 | 30 | 50 |  |
|  |  |  |  | 0.327 | 0.375 | 3.684 | 3.581 | < 0.001 | -21 | 21 | 47 |  |
| 0.002 | 0.007 | 27 | 0.001 | **0.013** | **0.081** | **4.602** | **4.413** | **< 0.001** | **-36** | **12** | **60** | Left middle frontal gyrus |
| 0.002 | 0.007 | 28 | 0.001 | **0.022** | **0.101** | **4.478** | **4.302** | **< 0.001** | **-12** | **-51** | **40** | Left precuneus |
|  |  |  |  | 0.074 | 0.127 | 4.171 | 4.027 | < 0.001 | -15 | -45 | 40 |  |
|  |  |  |  | 0.130 | 0.181 | 4.020 | 3.890 | < 0.001 | -9 | -57 | 43 |  |
| 0.228 | 0.324 | 4 | 0.152 | 0.060 | 0.127 | 4.225 | 4.076 | < 0.001 | 6 | 60 | 20 | Right medial superior frontal gyrus |
| 0.027 | 0.055 | 13 | 0.016 | 0.064 | 0.127 | 4.209 | 4.061 | < 0.001 | -30 | -93 | 10 | Left middle occipital gyrus |
|  |  |  |  | 0.618 | 0.713 | 3.373 | 3.293 | < 0.001 | -36 | -84 | 14 |  |
| 0.010 | 0.025 | 18 | 0.006 | 0.065 | 0.127 | 4.204 | 4.056 | < 0.001 | -45 | -51 | 24 | Left angular gyrus |
|  |  |  |  | 0.074 | 0.127 | 4.169 | 4.025 | < 0.001 | -45 | -57 | 24 |  |
| 0.052 | 0.088 | 10 | 0.031 | 0.149 | 0.181 | 3.973 | 3.847 | < 0.001 | 0 | -42 | 43 | Left cingulum |
|  |  |  |  | 0.152 | 0.181 | 3.968 | 3.842 | < 0.001 | 0 | -36 | 47 |  |
|  |  |  |  | 0.397 | 0.431 | 3.601 | 3.505 | < 0.001 | -9 | -42 | 43 |  |
| 0.405 | 0.474 | 2 | 0.306 | 0.156 | 0.181 | 3.958 | 3.834 | < 0.001 | -9 | -54 | 14 | Left precuneus |
| 0.174 | 0.273 | 5 | 0.113 | 0.380 | 0.429 | 3.620 | 3.523 | < 0.001 | -39 | -69 | 37 | Left middle occipital gyrus |
| 0.405 | 0.474 | 2 | 0.306 | 0.430 | 0.457 | 3.564 | 3.470 | < 0.001 | -36 | 24 | 40 | Left middle frontal gyrus |
| 0.553 | 0.474 | 1 | 0.474 | 0.462 | 0.481 | 3.530 | 3.439 | < 0.001 | -24 | 21 | 60 | Left superior frontal gyrus |
| 0.553 | 0.474 | 1 | 0.474 | 0.722 | 0.887 | 3.267 | 3.193 | < 0.001 | -3 | -27 | 47 | Left middle cingulum |
| 0.553 | 0.474 | 1 | 0.474 | 0.738 | 0.887 | 3.249 | 3.177 | < 0.001 | -12 | -27 | 40 | Left middle cingulum |
| 0.553 | 0.474 | 1 | 0.474 | 0.741 | 0.887 | 3.246 | 3.174 | < 0.001 | -45 | -63 | 50 | Left angular gyrus |
| 0.553 | 0.474 | 1 | 0.474 | 0.762 | 0.908 | 3.223 | 3.152 | < 0.001 | -3 | -24 | 40 | Left middle cingulum |
| 0.553 | 0.474 | 1 | 0.474 | 0.774 | 0.908 | 3.208 | 3.139 | < 0.001 | -48 | -54 | -16 | Left inferior temporal gyrus |
| 0.553 | 0.474 | 1 | 0.474 | 0.816 | 0.996 | 3.159 | 3.092 | < 0.001 | -9 | -30 | 43 | Left middle cingulum |

**Supplementary Table S13.**
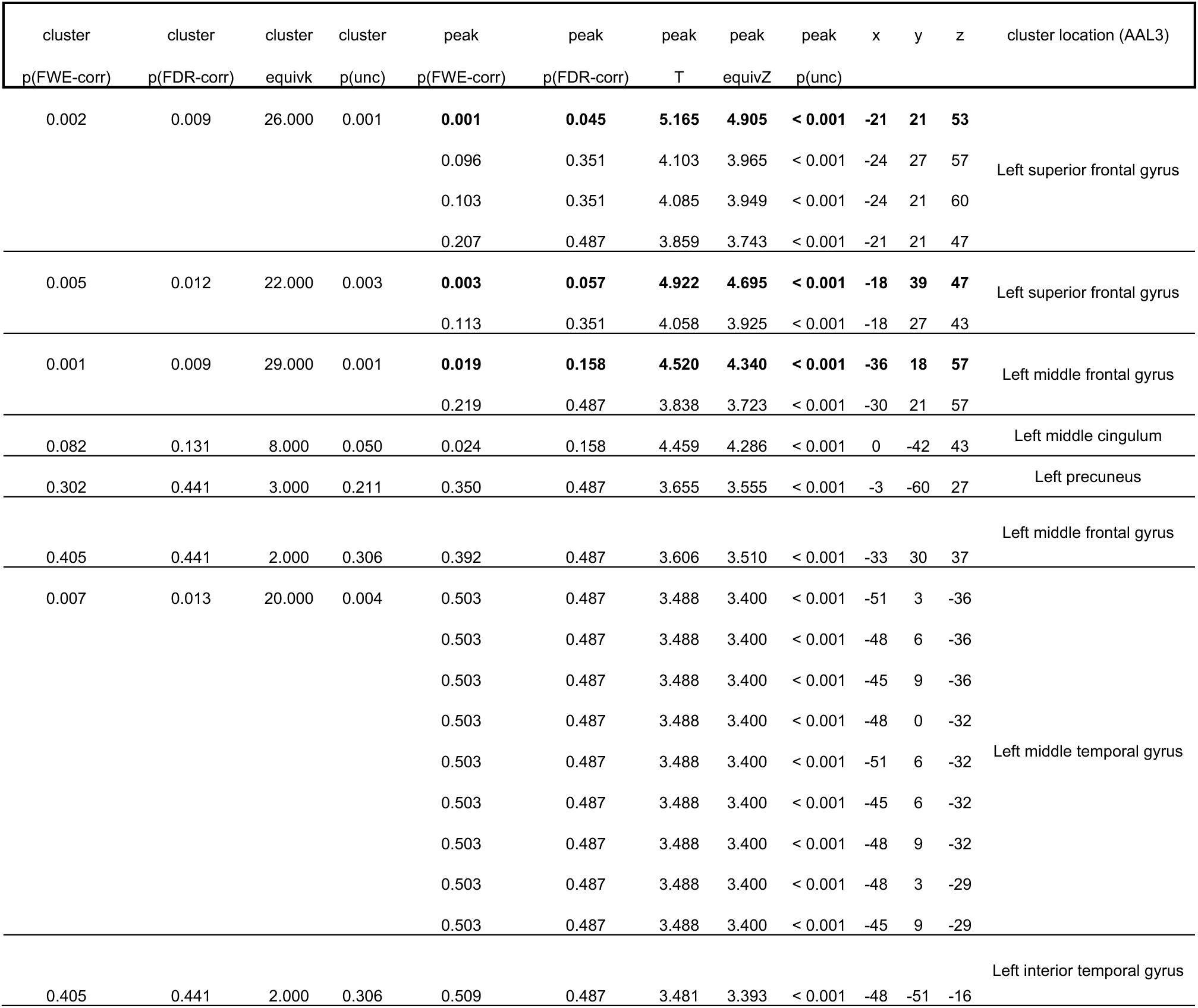
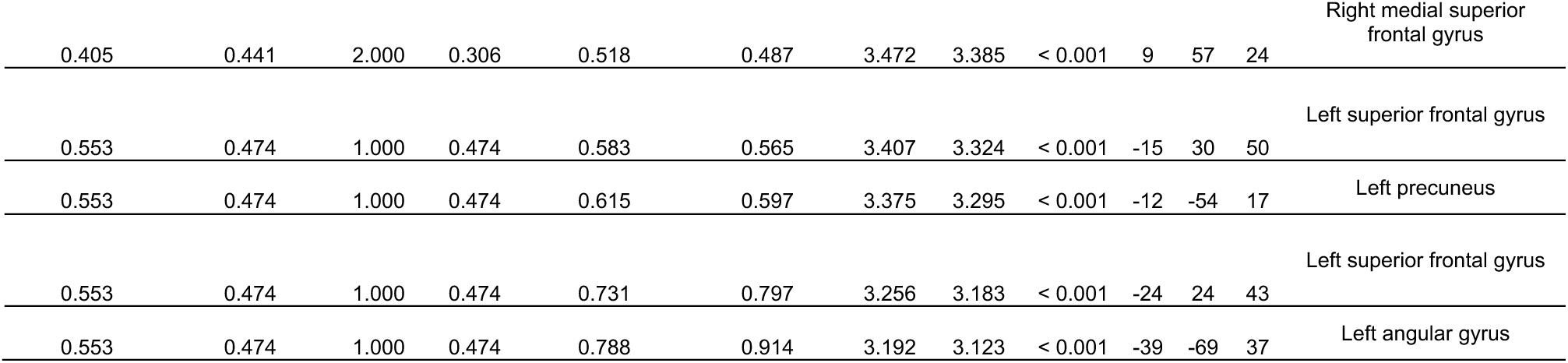
| RSA results for Morphologically Complex Existing Words. Coordinate in the standard MNI space of significant (p< 0.001 uncorrected, k>0) effect and cluster localization according to the AAL3 template^3^. Coordinates highlighted in bold are significant at a p< 0.05 FWE-corrected threshold. The term “k” indicates the total number of voxels in each cluster.

**Supplementary Table S14.**
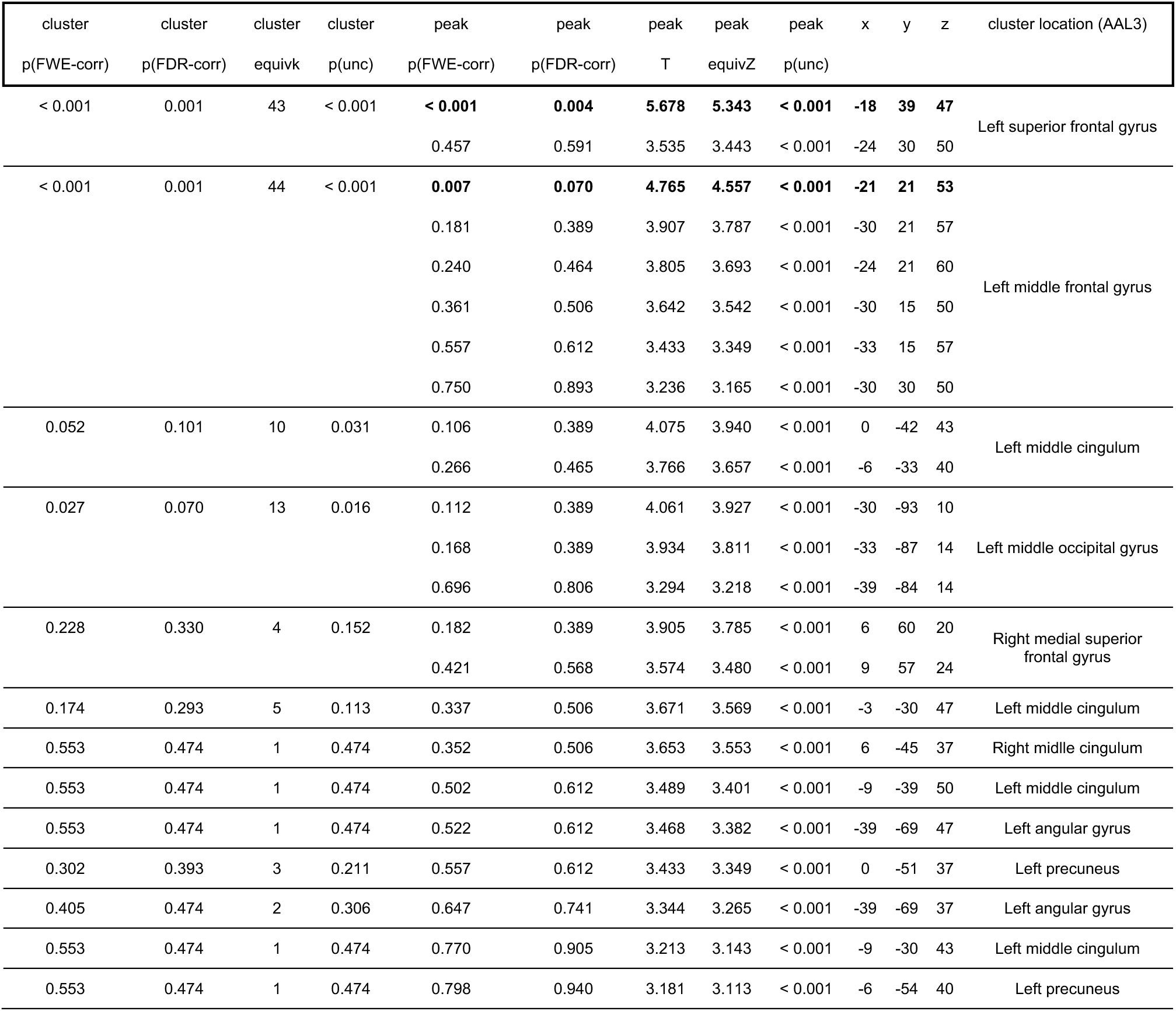
| Results of the Morphologically Complex > Morphologically Simple Existing Words contrast in RSA. Coordinates in the standard MNI space of significant (p< 0.001 uncorrected, k>0) effects and cluster localization according to the AAL3 template^3^. Coordinates highlighted in bold are significant at a p< 0.05 FWE-corrected threshold. The term “k” indicates the total number of voxels in each cluster.

**Supplementary Table S15.** | Results of the Morphologically Complex Novel Words > Morphologically Simple Novel Words contrast in RSA. Coordinates in the standard MNI space of significant (p< 0.001 uncorrected, k>0) effect and cluster localization according to the AAL3 template^3^. Coordinates highlighted in bold are significant at a p< 0.05 FWE-corrected threshold. The term “k” indicates the total number of voxels in each cluster.

| cluster<br>p(FWE-corr) | cluster<br>p(FDR-corr) | cluster<br>equivk | cluster<br>p(unc) | peak<br>p(FWE-corr) | peak<br>p(FDR-corr) | peak<br>T | peak<br>equivZ | peak<br>p(unc) | x | y | z | cluster location (AAL3) |
| --- | --- | --- | --- | --- | --- | --- | --- | --- | --- | --- | --- | --- |
| 0.033 | 0.070 | 12 | 0.020 | **0.049** | **0.160** | **4.275** | **4.120** | **< 0.001** | **-12** | **-51** | **40** | Left precuneus |
|  |  |  |  | 0.710 | 0.809 | 3.279 | 3.205 | < 0.001 | -9 | -57 | 40 |  |
| 0.010 | 0.041 | 18 | 0.006 | 0.055 | 0.160 | 4.247 | 4.096 | < 0.001 | -21 | 33 | 43 | Left superior frontal gyrus |
|  |  |  |  | 0.070 | 0.160 | 4.183 | 4.038 | < 0.001 | -27 | 36 | 43 |  |
| 0.052 | 0.073 | 10 | 0.031 | 0.320 | 0.562 | 3.692 | 3.589 | < 0.001 | -3 | -18 | 37 | Left middle cingulum |
| 0.105 | 0.114 | 7 | 0.065 | 0.379 | 0.562 | 3.620 | 3.523 | < 0.001 | -45 | -51 | 24 | Left angular gyrus |
|  |  |  |  | 0.532 | 0.744 | 3.458 | 3.372 | < 0.001 | -48 | -57 | 24 |  |
| 0.302 | 0.296 | 3 | 0.211 | 0.657 | 0.792 | 3.333 | 3.256 | < 0.001 | -36 | 12 | 60 | Left middle frontal gyrus |
| 0.553 | 0.474 | 1 | 0.474 | 0.659 | 0.792 | 3.331 | 3.253 | < 0.001 | -9 | -51 | 50 | Left precuneus |
| 0.553 | 0.474 | 1 | 0.474 | 0.778 | 0.886 | 3.204 | 3.135 | < 0.001 | -9 | -54 | 14 | Left precuneus |

## Notes

### Competing Interest Statement

The authors have declared no competing interest.

